# Extracellular vesicles associate with infectious geminiviral particles in the apoplast of infected plants

**DOI:** 10.64898/2026.05.22.727136

**Authors:** Pablo Morales-Martínez, Pepe Cana-Quijada, Benjamin L. Koch, Jerónimo Marulanda-Pulgarín, Rosa Lozano-Durán, Roger W. Innes, Eduardo R. Bejarano, Araceli G. Castillo

## Abstract

Plant viruses have evolved diverse strategies to facilitate their movement and survival within the host. Among them, geminiviruses co-opt host cellular machinery to replicate and disseminate. Traditionally, viral propagation has been associated with intercellular symplastic trafficking mediated by plasmodesmata and viral movement proteins. However, recent evidence demonstrated that the plant RNA virus turnip mosaic virus (TuMV) components are associated with extracellular vesicles (EVs). EVs are membrane-bound structures secreted to the extracellular space to potentially mediate several plant-pathogen interactions such as cross-kingdom RNA interference or the delivery of stress response proteins. In animals, EVs facilitate viral transmission both within the host and across species, but knowledge about their potential roles in plant viral infection is scarce.

In this study, we demonstrate that EVs isolated from geminivirus-infected plants contain complete viral genomes and both capsid and viral movement proteins. Furthermore, these EV fractions were demonstrated to be infectious when mechanically inoculated onto naïve plants. This discovery suggests that EVs may serve as alternative carriers for geminivirus components, enabling long-range transport or potentially modulating host immune responses, and highlights geminiviral capacity to transverse membrane boundaries, essential for circulative arbovirus propagation in their insect vectors.

**Significance Statement:** Viruses are obligate intracellular parasites that shape ecological communities and crucially challenge animal and plant health worldwide. Conversely to animal viruses, plant-infecting viruses rely on plasmodesmata to disseminate through their host and establish systemic infection. Nonetheless, most plant viruses are insect-transmitted whose ecological cycle relies on their insect vector spread. Thus, strategies to cross continuous membrane barriers are essential for their dissemination and may be potentially conserved in plant hosts. Our discovery that infectious viral entities are associated with EVs reveals an alternative pathway for geminiviral movement within plant hosts that could facilitate vector transmission, challenging our long-standing understanding of plant virus biology and expanding the current conception of plant viral pathology.

## Introduction

Successful viral infection requires the coordinated action of viral nucleic acids and proteins to hijack host cellular machinery, evade or suppress host defense responses, and support efficient replication in the infected cells. Beyond replication, the efficiency of viral propagation is highly dependent on the ability of the virus to spread from initially infected cells to neighboring tissues and ultimately reach the vascular system, which serves as the main route for long-distance dissemination. Viruses can achieve intercellular movement either by releasing the viral particles into the extracellular space or by direct cell–cell contact. In animal systems, the viral life cycle typically involves repeated crossing of cellular membranes during both egress from infected cells and entry into new target cells (2, 3). To overcome these membrane barriers, viruses have evolved diverse strategies that exploit host endocytic and exocytic pathways for efficient cell entry and exit (4, 5). Because plant cells are encased in the cell wall, they have evolved plasmodesmata to establish cytosolic continuity between cells. Plant viruses are known to co-opt plasmodesmata to symplastically spread through the plant host and establish systemic infection (6, 7). Nevertheless, the advent of pioneer methodologies in the analysis of plant extracellular landscape (8–10) has revealed the presence of viral elements in the plant extracellular space (apoplast), strengthening the possibility of apoplastic routes mediating viral dissemination, potentially dependent on extracellular vesicles (EVs) (1, 11).

EVs are membrane-bound compartments that carry diverse biological cargo (proteins, RNA species, lipids, and metabolites) secreted by the cell to the extracellular space and conserved in all three domains of life, *Bacteria, Archaea*, and *Eukarya* (12–14). EVs are hypothesized to mediate cell-to-cell communication, elicit defense responses and preserve cell homeostasis, potentially acting as protective carriers for long-distance transport (13, 15–17). The heterogeneity of EVs in cargo and functions is explained by their differences in size –ranging from 50 nm to 5 μm– and in their diverse biogenesis pathways: fusion of multivesicular bodies (MVBs) with the plasma membrane (PM), direct budding of microvesicles from the PM, or formation of large apoptotic bodies during cell death (14). In the plant kingdom, studies of EVs have lagged due to technical barriers such as the isolation of vesicles from the apoplast and the lack of molecular markers. However, in the last decade EVs from several plant species have been isolated and characterized, including *Arabidopsis thaliana* (from this point on Arabidopsis) (18–20) and other relevant crops such as sorghum, tobacco, alfalfa, and rice (21–25). Upon abiotic or biotic stresses, plants upregulate EV secretion, commonly enriched in defense-related proteins, such as the SYNTAXIN OF PLANTS 121 (SYP121) (also called PENETRATION 1, PEN1) and the tetraspanin TET8 (8, 19). These markers label distinct populations of plant EVs that can be separated by differential ultracentrifugation (9, 26).

Despite the rapidly growing understanding of EV-mediated communication in plants, very little is known about the potential involvement of EVs in plant-virus interactions. This gap is particularly striking given the well-documented roles of EVs in animal virus biology, where both enveloped and non-enveloped viruses exploit EVs to enhance replication, suppress immune responses, protect from proteases or nucleases, and promote *en bloc* insect vector spread (reviewed in 22–26). Nonetheless, in 2019 Movahed et al. (1) demonstrated for the first time the release of the potyvirus turnip mosaic virus (TuMV) proteins and RNAs to the extracellular space associated with EVs in infected Arabidopsis plants. Moreover, virions from the potexvirus Potato virus X (PVX) have been detected in the apoplast of infected hosts (11), opening the possibility of the existence of noncanonical viral movement pathways apart from the plasmodesmata-mediated transport.

To further explore this potential transport, we screened plants infected with geminiviruses, mainly focusing on the begomovirus cabbage leaf curl virus (CabLCV). Begomoviruses are plant-infecting DNA viruses with single-stranded circular genomes encapsidated in twinned icosahedral particles. These viruses are exclusively transmitted by the whitefly *Bemisia tabaci* and cause major crop losses worldwide, particularly in tropical and subtropical regions (32–34). Begomoviruses are classified as monopartite or bipartite according to their genomic organization, either in a single component or in two molecules named DNA A and DNA B, respectively. DNA A, which encodes the coat protein (CP), resembles the monopartite genome, while DNA-B encodes two proteins required for viral movement in the plant: the nuclear shuttle protein (NSP), and the movement protein (MP), which cooperatively move the viral genome from its site of replication in the nucleus to the cytoplasm and into adjacent plant cells (35). In bipartite begomoviruses, CP, NSP and MP function cooperatively to direct this viral DNA movement, while in monopartite begomoviruses, some of these functions are carried out by CP assisted by other viral proteins such as V2 and C5 (35). Although the mechanisms by which viral complexes reach the plasma membrane remain unresolved, available evidence indicates that trafficking depends on interactions between viral complexes, composed of movement proteins and viral DNA, and host factors. Accordingly, MP interaction with synaptotagmin A (SYTA) supports a model in which geminiviral complexes are targeted to plasmodesmata via an endocytic recycling pathway (36, 37). Alternatively, a stromule network connecting the nucleus to plasmodesmata has been proposed, with cpHSC70-1 acting as a docking platform for viral complexes (38, 39). Finally, microtubule-associated factors (Pin4, SCD2) and actin-dependent ER trafficking appear to further contribute to MP delivery to plasmodesmata (40, 41). Although it is widely accepted that begomovirus intercellular movement occurs through plasmodesmata, important questions remain unresolved, including the precise form of viral DNA exported, molecular details of viral engagement of host export machinery, and the contribution of cytoskeletal and endomembrane routes. Furthermore, the biological relevance of the vesicle trafficking system in geminiviral movement has been recently demonstrated, as virus induced gene silencing (VIGS) of the essential vesicle trafficking genes *δ-COP, ARF1, CHC1* and *CHC2* in *Nicotiana benthamiana* completely abolished systemic viral accumulation (42, 43).

Here, we demonstrate that i) begomovirus infection induces the accumulation of EVs in the apoplast of Arabidopsis; ii) viral DNA together with viral proteins involved in encapsidation and movement are released into the extracellular space in association with EVs; and iii) these nucleic acids are protected within vesicular structures. Importantly, we also show that apoplastic extracts from infected Arabidopsis plants are sufficient to transmit infection to healthy plants, indicating that EVs can provide geminivirus genome protection and horizontal transmission.

## Results

### Geminivirus infection induces the release of EVs enriched in stress-response proteins to the host plant apoplast

To evaluate the impact of begomovirus infection in EV release to the apoplast, apoplastic wash fluid (AWF) was collected from both CabLCV-infected and non-infected RFP-PEN1 × TET8-GFP plants at 15 days post-inoculation (dpi). Then, EV fractions were collected after a 40 000 ×g ultracentrifugation (P40) and visualized through Confocal Laser Scanning Microscopy (CLSM) or Transmission Electron Microscopy (TEM) to confirm the isolation of EVs. CLSM detected RFP-PEN1 and TET8-GFP signals in distinct, non-overlapping EV populations, as previously reported (9, 10) (Fig.1A), and TEM revealed typical cup-shaped vesicles ranging from 100 nm to 300 nm, consistent with Arabidopsis EV morphology (8). Additionally, polysaccharide filaments from the cell wall were also observed as in (9) (Fig.1B). Furthermore, detection by Western blot of EV marker proteins RFP-PEN1, TET8-GFP and PATELLIN 1 (PATL1) confirmed the extraction of EVs in P40 samples, while the SYP61 signal, an intracellular marker protein associated with TGN (8), was only observed in cell lysate samples but not in P40 fractions (Fig.1C) implying that P40 fractions tested were devoid of intracellular contaminations. Interestingly, immunodetection of EV markers showed a clear increase in P40 fractions from CabLCV-infected plants (Fig.1C; Table S1). These results were validated by Nanoparticle tracking analysis (NTA), showing that AWF from CabLCV-infected plants is significantly enriched in EVs compared to the non-infected condition (Fig.1D). Additionally, a marked increase in both PEN1- and TET8-positive EV populations was detected by CLSM in P40 pellets isolated from CabLCV-infected plants compared to the non-infected ones (Fig.1E). Taken together, these results support the notion that EV secretion is induced during CabLCV infection.

**Fig. 1.**
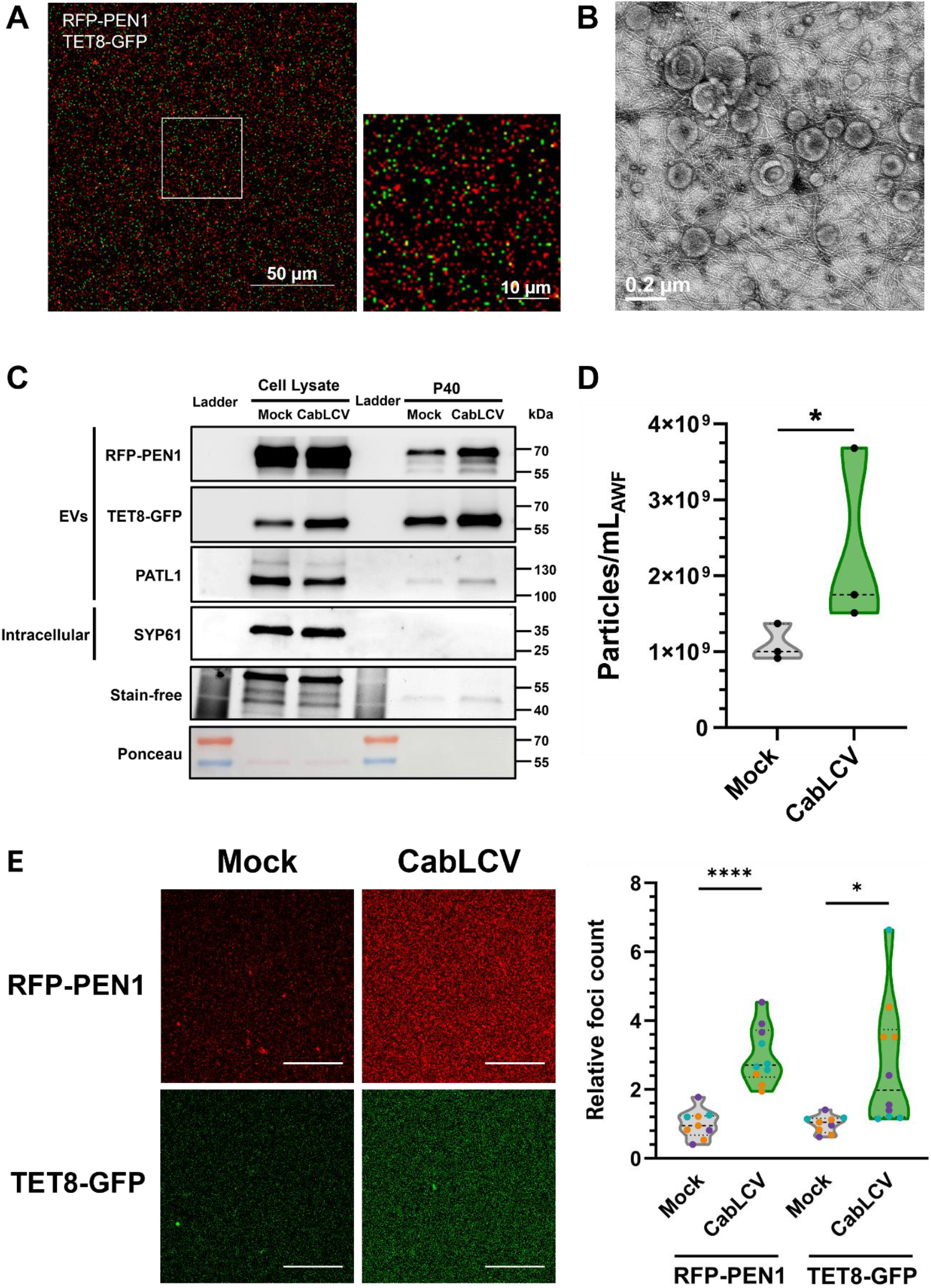
CabLCV infection promotes EV release to the apoplast of *Arabidopsis thaliana* plants. EV isolation procedures described in Rutter & Innes, 2017 (8) and Rutter et al. 2017 (18) allowed successful extraction of EVs confirmed by **(A)** the detection of fluorescent signal from EV-marker proteins in P40 pellets from RFP-PEN1 × TET8-GFP *Arabidopsis thaliana* transgenic lines using Confocal Laser Scanning Microscopy (CLSM), **(B)** Transmission Electron Microscopy (TEM) visualization of cup-shaped structures ranging 100 nm – 300 nm and **(C)** immunodetection of EV-marker proteins RFP-PEN1, TET8-GFP and PATL1. A qualitative increase in EV-marker proteins was observed for CabLCV-infected plants compared to mock-treated plants in two biological replicates and further confirmed by **(D)** Nanoparticle Tracker Analysis (NTA), showing a significant increase in the amount of nanoparticles per mL of AWF in P40 fractions isolated from CabLCV-infected plants compared to mock-treated plants in three biological replicates. **(E)** Furthermore, the enhanced release of EVs to AWF in CablCV-infected plants occurs simultaneously for both RFP-PEN1 and TET8-GFP EV-marker protein fluorescent signals by CLSM. Bars in panel E represent 50 μm. RFP-PEN1 and TET8-GFP EV foci quantification was relativized to the non-infected control (Mock, set to 1). Asterisks represent statistically significant difference with the Mock control according to a ratio paired Student’s t-test (****, P-value < 0.0001; *, P-value < 0.05). The experiment was conducted in three independent replicates, yielding consistent results and distinguished by dot colours (purple, orange and blue).

### Apoplast from infected plants carry geminiviral genomes associated with EVs

The association of begomovirus genomes with EVs in the apoplast was subsequently addressed. AWF and the corresponding P40 vesicle-enriched fractions were isolated from CabLCV-infected and mock-inoculated plants and the presence of viral DNA was assessed using PCR. Geminiviral DNA was consistently detected in the AWF and P40 fractions from infected plants, whereas no amplification was observed in samples derived from mock-inoculated controls (Fig. 2A). As an internal control, the presence of the actin gene was examined to assess potential nuclear contamination in the apoplastic preparations. Actin sequences were not amplified from either the AWF or P40 fractions, while they were readily detected in leaf cell lysate DNA extractions. These results confirm that the viral DNA detected in apoplastic and vesicle-enriched fractions does not arise from nuclear contamination (Fig. 2A). We then examined whether the viral DNA detected in P40 fractions was freely exposed or physically protected within vesicular structures. To this end, a DNase protection assay was performed. This approach relies on the principle that naked DNA is readily degraded by nucleases, whereas DNA enclosed within compartments remain resistant to enzymatic digestion. Viral DNA was consistently detected in CabLCV-infected P40 fractions, even when treated with DNase, indicating that the viral DNA present in EV fractions is protected from nuclease activity (Fig. 2B, left).

**Fig. 2.**
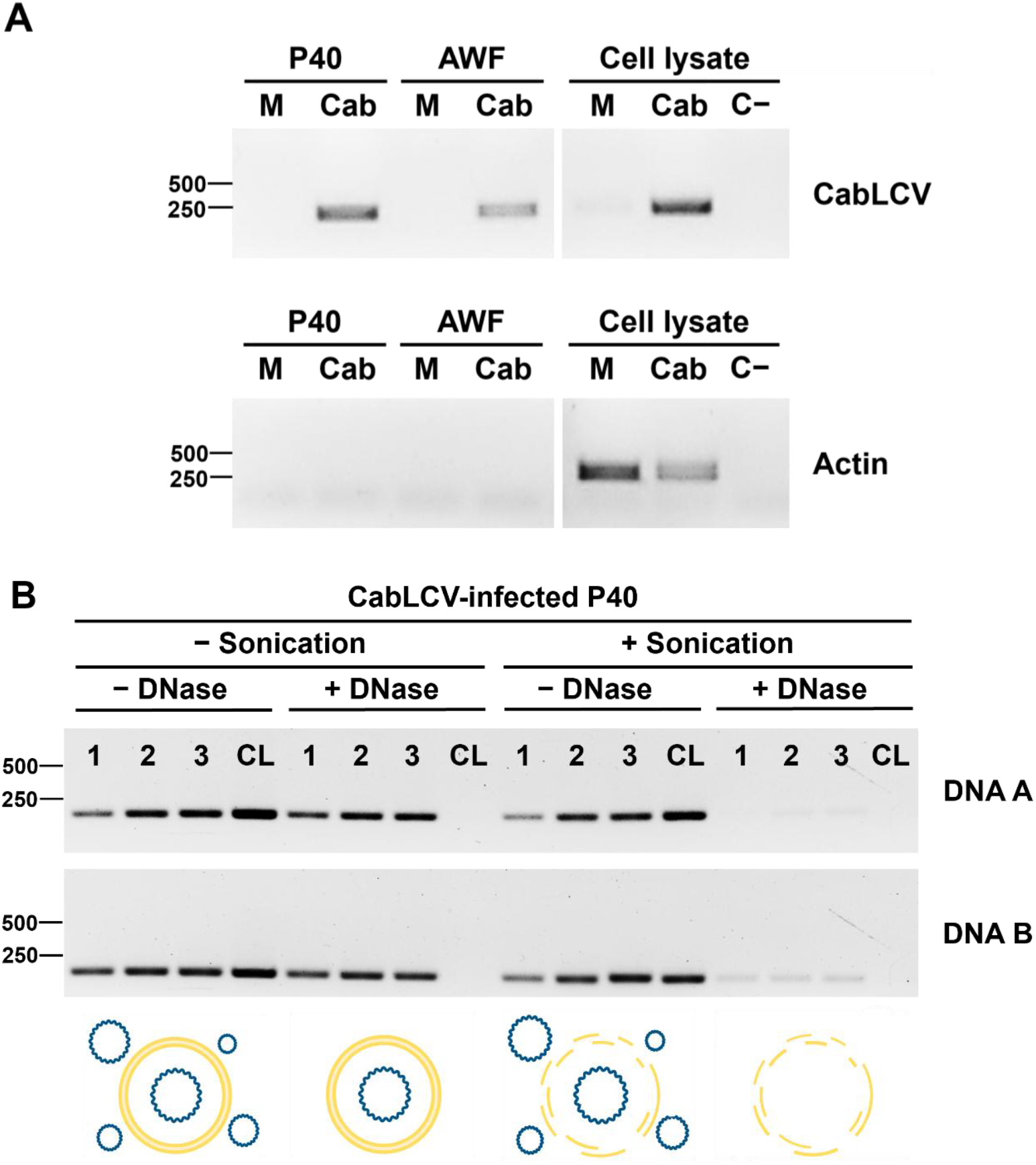
CabLCV DNA sequences are detectable and protected in P40 fractions from infected plants. **(A)** CabLCV DNA sequences from AL3 gene (DNA A) were amplified by PCR in AWF and P40 fractions isolated during the EV isolation protocol. The absence of amplification of actin gene suggest that CabLCV DNA amplification is not derived from nuclear contamination. **M**: Samples from mock-treated plants. **Cab**: Samples from CabLCV-infected plants. **C™**: DNA-free negative control **(B)** Incubation of P40 fractions from CabLCV-infected plants with DNase only prevented geminiviral DNA amplification when previously sonicated, suggesting that viral DNA sequences are confined within protective structures. PCR amplification targeted genes AL3 for DNA A and BC1 for DNA B. **1-3**: independent P40 fractions from the same batch of AWF. **CL**: Cell lysate DNA extraction from CabLCV-infected plants. Experiments were reproduced twice with similar results.

To further test whether the observed protection of viral DNA was due to vesicle encapsulation, we applied sonication and heat shock to P40 fractions to disrupt the vesicle prior to DNase treatment. We confirmed that, in treated P40 samples, viral DNA was only detected in the aliquots not treated with DNase. In contrast, no amplification was observed in sonicated samples following DNase digestion, demonstrating that vesicle disruption renders the viral genome accessible to nuclease degradation (Fig. 2B). PCR analyses were performed using specific oligonucleotides targeting both components of the CabLCV genome (DNA A and DNA B). Consistent results were obtained for each component, indicating that both genomic components are equally protected from DNase activity (Fig. 2B). To verify that sonication and heat treatment effectively compromise vesicle integrity, we employed an approach similar to that described in Zand Karimi et al., 2022 (44). P40 pellets were subjected to ultrasounds and thermal shock, followed by treatment with trypsin, and the level of the EV-associated marker protein RFP–PEN1 was assessed by Western blot. The RFP immunoblot signal, which remains largely protected from proteolysis under non-disruptive conditions (decreasing slightly from 1 to 0.61), showed a ∼2–3-fold reduction (from 1 to 0.23) when P40 fractions were sonicated prior to trypsin digestion, consistent with increased exposure of EV luminal contents (Fig. S1).

The activity of the DNase enzyme was verified by applying the same treatment to cell lysate DNA extractions from CabLCV-infected plants (CL). In this case, the enzyme efficiently degraded the DNA, confirming that the apparent protection of viral DNA in infected samples was not due to loss of enzymatic activity (Fig. 2B). Taken together, these data demonstrate that CabLCV DNA present in P40 fractions is physically protected from nuclease digestion, indicating that the loss of viral DNA protection after sonication and heat shock results from vesicle disruption rather than from indirect enzymatic effects. Importantly, this protective association was not limited to CabLCV: comparable results were obtained from apoplastic fractions of Arabidopsis plants infected with the geminivirus beet curly top virus (BCTV, genus *Curtovirus*) (Fig. S2).

The presence of complete circular CabLCV genomes in extracellular vesicle fractions was evaluated using rolling circle amplification (RCA), a technique that detects circular DNA molecules by generating high–molecular-weight concatemeric products. To characterize the amplified products and confirm the amplification of full-length viral genomes, RCA products were digested with restriction enzymes that cleave both CabLCV genome molecules at a single site, generating linear fragments of the expected size of approximately 2.5 kb for both DNA A and DNA B (Fig. 3A). Strikingly, RCA performed on P40 fractions from CabLCV-infected plants yielded high-molecular-weight products that, upon digestion with *Sac*I, resolved into distinct monomeric bands of ∼2.5 kb. Identical results were obtained using cell lysate DNA extracts from CabLCV-infected plants, which served as positive controls. In contrast, no amplification or restriction fragments were detected in P40 fractions from mock-inoculated plants, confirming the specificity for the viral signal (Fig. 3B).To further validate the identity of the amplified molecules and discriminate between DNA A and DNA B components, RCA products were digested with enzymes that differentially cleave the two viral genomes (Fig. 3C). Digestion with *Pvu*II, which cuts only once in DNA B but not in DNA A, released a single ∼2.5 kb band, confirming the presence of DNA B. By contrast, digestion with *Bsi*I, which cuts once in DNA B and twice in DNA A, produced three fragments corresponding to the expected sizes for both components: a ∼2.5 kb band from DNA B and two bands of ∼1.8 and ∼0.8 kb from DNA A. These results establish that both viral genomic components are present as intact circular molecules in the P40 fractions from infected plants. The molecular identity of the ∼2.5 kb fragment released by *Sac*I digestion was finally confirmed by cloning and sequencing (Fig. S3), verifying its correspondence to CabLCV DNA A and DNA B. Together, these findings demonstrate that extracellular vesicle-enriched fractions from infected plants contain complete, circular DNA-A and DNA-B CabLCV genomes. Similar RCA results were obtained in P40 fractions isolated from *N. benthamiana* infected and mock-inoculated plants (Fig. S4), further supporting that the accumulation of complete, circular CabLCV genomes within extracellular vesicle–enriched fractions is conserved across plant hosts.

**Fig. 3.**
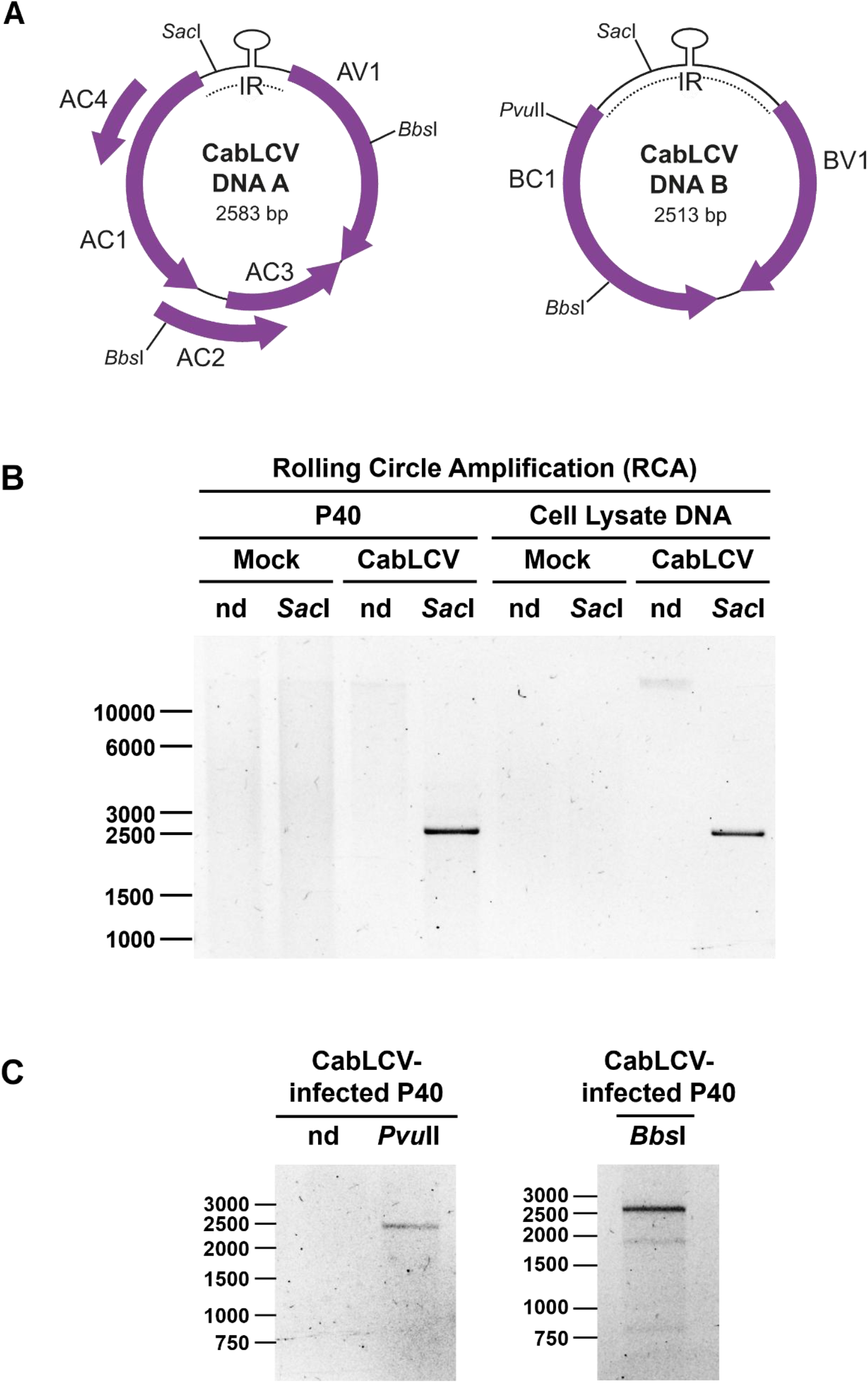
Complete geminiviral genomes were amplified in P40 fractions from CabLCV-infected plants. **(A)** CabLCV DNA A and DNA B genomic component maps indicating target sites of selected endonucleases used to address the identity of RCA products. **(B)** Rolling Circle Amplification (RCA) allowed detection of complete circular CabLCV genomes in P40 fractions. RCA products were linearized using single-cutter endonucleases, releasing monomers of ∼2.5 kb. RCA using cell lysate DNA extractions as templates were included as positive controls. Experiments were reproduced in three independent biological replicates with similar results. **(C)** Restriction analysis of RCA products once incubated with selected endonucleases yielding differential patterns for DNA A (non-cut with *Pvu*II and scised in two fragments of 1804 bp and 779 bp by *Bbs*I) and DNA B (linearized with *Pvu*II and *Bbs*I). **nd:** non-digested. ***Sac*I:** *Sac*I-digested. ***PvuI*I:** *Pvu*II-digested. ***Bbs*I**: *Bbs*I-digested.

### Movement-related CabLCV proteins are detected in the P40 fraction of infected Arabidopsis plants

To investigate whether additional viral components are present in the vesicle-enriched P40 fractions, we performed a proteomic analysis of these samples. Proteins isolated from P40 fractions of CabLCV-infected plants were analyzed by label-free quantitative mass spectrometry (MS) (Table 1). In total, 1,989 proteins were confidently identified with high peptide coverage. Among CabLCV viral proteins, three were identified with global peptide-spectrum matches equivalent to 316, 236 and 3, corresponding respectively to the coat protein CP, and the two proteins encoded by DNA B, the nuclear shuttle protein NSP (BV1) and the movement protein MP (BC1), essential for intracellular and systemic movement of bipartite geminiviruses. In addition to viral proteins, numerous host proteins typically associated with EVs were detected, including established EV markers such as the PEN1 (AT3G11820), TET8 (AT2G23810), and PATL1 (AT1G72150). Furthermore, proteins involved in membrane trafficking and stress responses, such as RIN4 and other defence-associated components, were also represented, whereas already described intracellular and plasma membrane marker proteins such as SYP61 (trans-Golgi Network), ARA6 (MVB/Late Endosome) or LTI6b (PM) were not found (8), supporting the absence of intracellular contamination in our samples.

**Table 1.** P40 fractions from CabLCV-infected plants contain geminiviral proteins detectable by LFQ-MS. Selection of viral proteins and EV-marker proteins found in proteomic analysis of P40 fractions from CabLCV-infected plants collected in three independent biological replicates. intracellular markers previously described to be absent in P40 fractions (SYP61 for TGN, ARA6 for MVB/LE or LTI6b for PM) (8) were not detected. Values for sum of posterior error probability (Sum PEP Score), global peptide-spectrum matches (Global PSM), global peptides (Global Pepts), global unique peptides (Global Uniq. Pepts) and abundance are shown for each protein enlisted.

|  | Accession<br>TAIR | Protein Name | Sum PEP<br>Score | # Global<br>PSMs | # Global<br>Pepts | # Global<br>Uniq. Pepts | Abundance |
| --- | --- | --- | --- | --- | --- | --- | --- |
| **Viral Proteins** | Not<br>applicable | **AV1 (CP)** | 126.576 | 316 | 10 | 10 | 4.05E+07 |
|  |  | **BC1 (MP)** | 5.689 | 3 | 1 | 1 | 8.08E+05 |
|  |  | **BV1 (NSP)** | 102.423 | 236 | 5 | 5 | 3.20E+07 |
| **EV marker**<br>**proteins** | AT3G11820 | **Syntaxin-121 (PEN1)** | 665.975 | 4908 | 26 | 26 | 6.93E+08 |
|  | AT2G23810 | **Tetraspanin-8 (TET8)** | 24.922 | 411 | 4 | 4 | 5.23E+07 |
|  | AT1G72150 | **Patellin-1 (PATL1)** | 334.128 | 1247 | 28 | 26 | 1.01E+08 |
|  | AT3G25070 | **RPM1-interacting**<br>**protein 4 (RIN4)** | 30.401 | 42 | 2 | 2 | 3.09E+06 |

Approximately90% of the proteins identified in (185 of 204) previously reported in published Arabidopsis thaliana EV proteomes (8) (185/204) overlap with accessions found in thewere also detected in our CabLCV-P40 samples, further supporting the EV origin of the P40 fraction. Additionally, GO enrichment analysis confirms the presence of strong signal for apoplast-related categories such as “*secretory vesicle*”, “*apoplast*” or “*cell wall*” (Table S2). These findings demonstrate that the vesicle-enriched P40 fractions contain not only complete CabLCV DNA molecules, but also viral proteins required for genome encapsidation and movement, together with host proteins characteristic of extracellular vesicle biogenesis and function. These results further validate the vesicular origin of the P40 fraction and highlight the complex molecular composition of infection-induced EVs.

### The extracellular vesicle–associated geminiviral components are infectious

To assess whether the extracellular vesicle–associated viral components are infectious, P40 fractions from CabLCV-infected plants were used to mechanically inoculate Arabidopsis leaves. These P40 pellets were confirmed by PCR analysis to harbour non-nuclear CabLCV sequences as no amplification was detected for actin oligonucleotides (Fig. 4A). Inoculated plants developed characteristic CabLCV symptoms at 28 dpi (Fig. 4B) and tested positive for viral DNA by RCA amplification of full-length genomes and sequencing confirmation (Fig. 4C). These findings demonstrate that EVs released during infection contain viral elements capable of initiating a complete infection cycle.

**Fig. 4.**
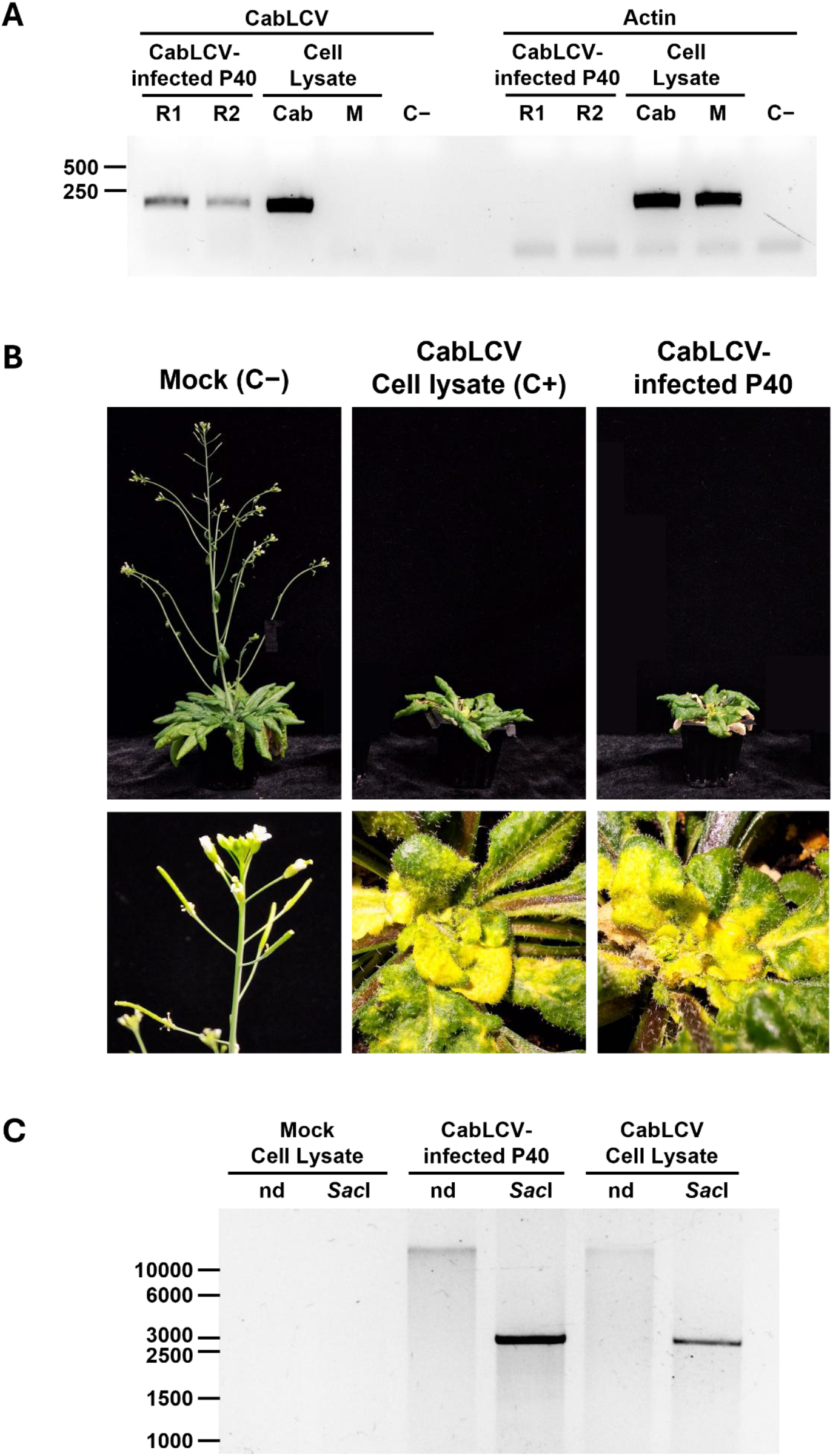
P40 fractions from CabLCV-infected plants are infectious in naive *Arabidopsis thaliana* plants. **(A)** P40 fractions used for mechanical inoculation were confirmed to harbour CabLCV sequences not deriving from the nucleus as no actin amplification was achieved. P40 samples from two independent biological replicates were included (R1-R2). **M**: Cell lysate DNA from mock-treated plants. **Cab**: Cell lysate DNA from CabLCV-infected plants. **C™**: DNA-free negative control. **(B)** Naive *Arabidopsis thaliana* plants mechanically inoculated with P40 fractions isolated from CabLCV-infected plants at 21 days after agroinoculation developed typical CabLCV symptoms at 28 dpi. Buffer-alone inoculation and cell lysate extraction from CabLCV-infected plants were respectively used as negative (Mock) and positive (CabLCV) controls. Experiments were reproduced in three independent biological replicates with similar results. **(C)** DNA isolated from P40-inoculated plants was assayed by RCA confirming the presence of CabLCV viral genomes.

Collectively, our results suggest that these infectious viral elements found in P40 pellets may harbour viral genomes associated with CP and movement proteins NSP/MP. However, whether any of these nucleoproteic complexes form and are loaded to EVs remains unclear. As virions constitute the main infectious form of viruses outside cells and CP is essential for capsid formation, we asked whether CP was required for viral release into the apoplast. To address this question, plants were infected with a CabLCV^ΔCP^ mutant, and P40 fractions were analyzed for the presence of viral DNA. Strikingly, full-length viral genomes were still detectable in the P40 fractions of plants infected with the CP mutant (Fig.5), confirming that the presence of CabLCV DNA in P40 fractions does not require CP and therefore cannot be exclusively explained by the formation of canonical virions. Instead, viral DNA may be encapsulated or stabilized by EVs or associated host-derived structures, providing alternative means of extracellular protection and transport that is independent of CP-mediated virion assembly.

**Fig. 5.**
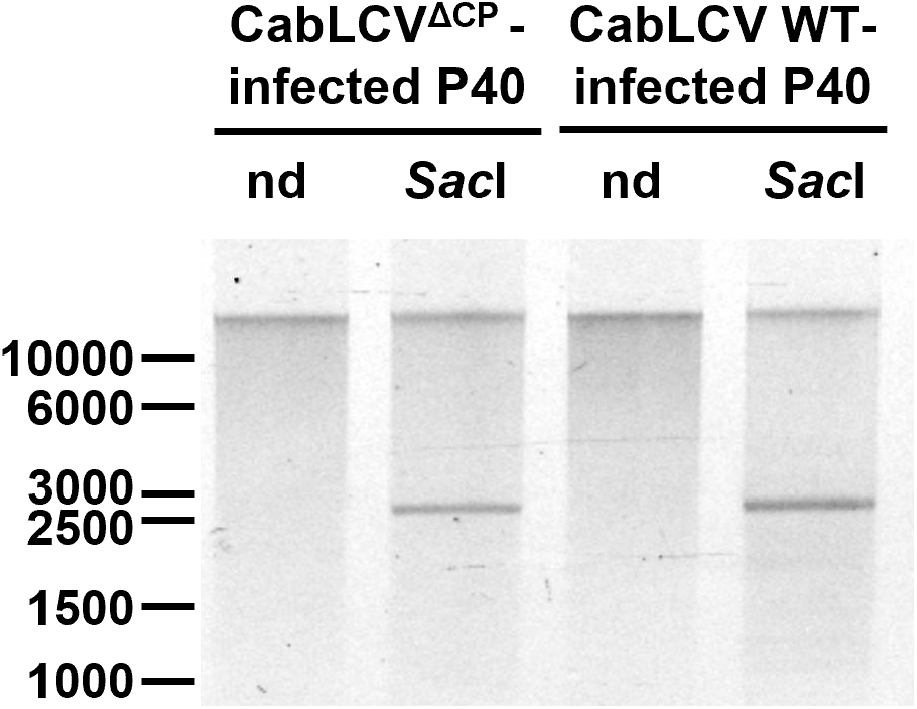
Viral genomes are detected in P40 fractions from CabLCV^ΔCP^-infected plants. Rolling Circle Amplification (RCA) allowed detection of complete circular CabLCV genomes in P40 fractions both from wild-type CabLCV and mutant CabLCV^ΔCP^ infected plants. RCA products were linearized using single-cutter endonucleases. RCA using cell lysate DNA extractions as templates were included as positive controls. Experiments were reproduced twice with similar results. **nd:** non-digested. ***Sac*I:** *Sac*I-digested.

CP from CabLCV contributes to but is not essential for plant systemic infection. At 21 dpi with CabLCV^ΔCP^, viral DNA quantification revealed an approximately 30-fold and 9-fold decrease in DNA A and DNA B accumulation in systemic leaves, respectively, in the mutant compared with the wild type (Table S3). To determine whether viral DNA accumulated from CabLCV^ΔCP^-infected plants could establish systemic infection upon mechanical inoculation, extracts from infected plants were used as inoculum. Because viral DNA accumulation differed between WT- and ΔCP-infected plants, wild-type extracts were diluted to equalize viral DNA concentrations prior to inoculation. None of the plants inoculated with ΔCP extracts (0/65) developed infection symptoms, whereas the diluted wild-type inoculum achieved a 22.7% infection rate (Table 2). These results were further confirmed by PCR, as none of the randomly selected plants inoculated with CabLCV^ΔCP^ extracts amplified viral sequences, in contrast to plants inoculated with the wild-type inoculum (Fig. S5). These findings indicate that CP presence is essential for systemic infection of CabLCV following mechanical inoculation and suggest that mechanical inoculation with extracellular vesicle– associated CabLCV^ΔCP^ virus would be ineffective. Together, these results establish that apoplastic EVs act as vehicles for infectious geminiviral DNA, with the coat protein playing a pivotal role in enabling their stability and transmission.

**Table 2.**
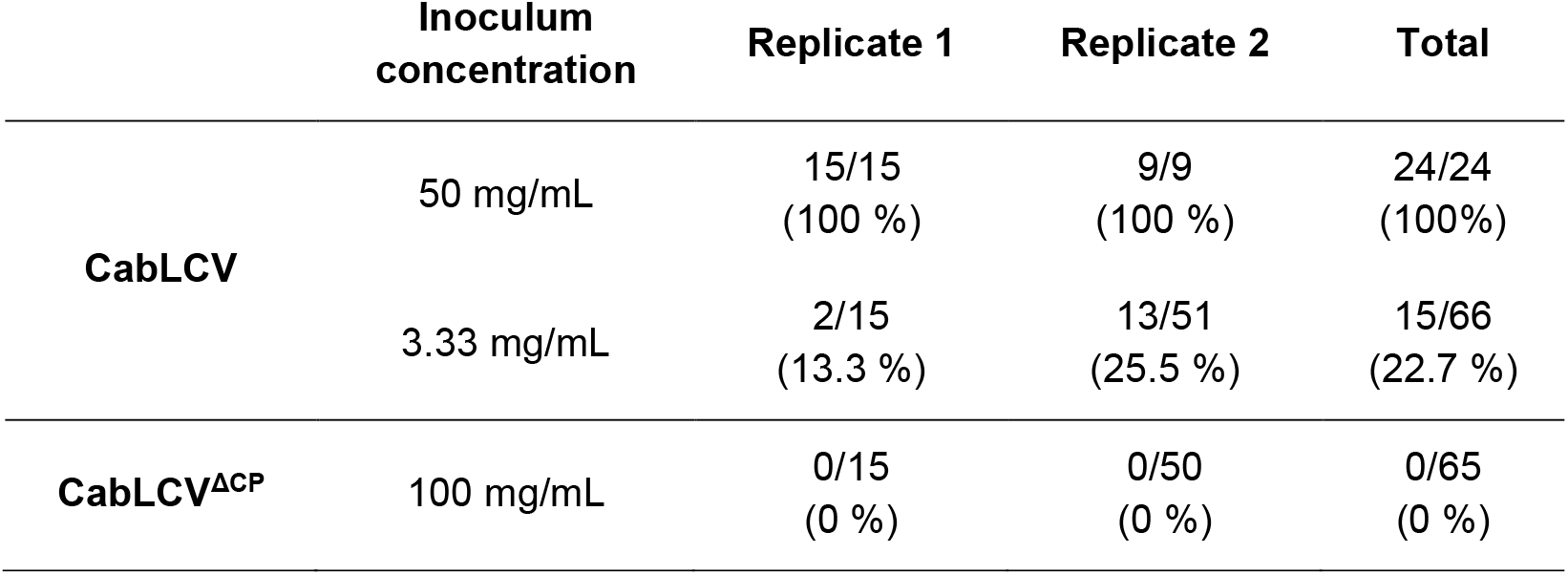
CP is essential for CabLCV systemic infection following mechanical inoculation. Naive *Arabidopsis thaliana* plants mechanically inoculated with extracts from plants infected either with CabLCV or mutant CabLCV^ΔCP^ and containing equivalent doses of DNA A component developed only viral symptoms when rubbed using wild-type CabLCV. Plants inoculated with standard amount of ground tissue (50 mg/mL) were used as positive controls.

## Discussion

This study provides direct evidence that EVs isolated from geminivirus-infected plants contain complete viral genomes together with key viral proteins involved in encapsidation and viral movement. The combination of DNase and trypsin protection assays, together with sonication and heat shock treatment, strongly supports that viral DNA is tightly associated with intact vesicles, though high-resolution single-vesicle imaging and cryo-electron microscopy will be required to resolve the precise architecture of these viral cargoes and to distinguish true intraluminal encapsidation from surface association or association with other plant extracellular particles or cell wall filaments.

Although the origin and biogenesis of EVs containing geminiviral infectious particles remain unresolved, multiple lines of evidence point to the endosomal pathway and MVBs as a likely source. First, during infection, geminiviral DNA associates with coat protein CP and movement proteins (NSP and MP) in endosomal compartments that may give rise to MVBs, a major source of EVs and cellular structures implicated in intra- and intercellular viral movement (1, 45). Second, EV-associated markers such as TET8 and PEN1, detected in EVs from infected plants, have also been localized to MVB-like compartments (10). Third, reducing COPI activity, a complex that has been implicated in the formation of ILVs in MVBs (46), severely impairs geminiviral spread without affecting replication (43). Future studies will be required to define the cellular origin of these vesicles and the mechanisms underlying selective viral cargo loading. In particular, it remains to be determined whether geminivirus movement proteins only engage components of the MVB pathway or alternative secretion routes are involved, such as the ones involving EXPO (exocyst-positive organelle) secretion (10, 13).

### Biological relevance of the viral association to EVs in plants

Although the biological relevance of infectious viral particles associated with EVs remains to be fully established, it is plausible—based on the results presented here and insights from animal systems—that such associations could contribute to protection from degradation or immune detection, facilitate intercellular transfer, or broaden mechanisms of cell entry (28, 30).

Protection from extracellular nucleases and environmental stress is a key functional consequence of EV encapsulation. The protection assays presented in this study demonstrate that viral DNA is shielded from the degradation by DNases, likely enhancing the stability of viral genomes in the apoplast, where free nucleic acids would otherwise be rapidly degraded. On the other hand, EVs are increasingly recognized as active regulators of immune signalling and intercellular communication rather than passive carriers. In plants, pathogen challenge induces EV biogenesis, while salicylic acid signalling promotes the selective loading of antimicrobial cargo and mobile signals that prime neighbouring cells, thereby amplifying immunity across tissues (47, 48). Our data show that geminivirus infection similarly activates EV production. This response may have opposing consequences: it could favour infection, as widely observed in animal systems where viruses exploit exosomal pathways to enhance their spread, or it could restrict infection by potentiating systemic acquired resistance (SAR). Dissecting these alternatives will require functional analyses of virus-associated EV populations and mutants impaired in EV biogenesis.

Although plant viruses are obligate intracellular pathogens, accumulating evidence indicates that they can activate pattern-triggered immunity (PTI), likely through direct or indirect recognition of viral components by plasma membrane-localized receptor-like kinases (RLKs) (49– 52). This process appears analogous to the perception of extracellular pathogen-associated molecular patterns (PAMPs) during bacterial, fungal, or arthropod attack. In this context, the presence of viral DNA and proteins in the apoplast provides a plausible mechanistic basis for such recognition. Conversely, the encapsulation of viral components within EVs may facilitate immune evasion by limiting their accessibility to RLKs, thereby reducing the likelihood of PAMP perception.

### EVs and viral movement in plants

Although the presence of viral particles in the plant apoplast has been previously reported for two positive-sense RNA viruses (the potyvirus Turnip mosaic virus (TuMV), whose components are secreted in vesicles derived from MVBs (1), and the potexvirus Potato virus X (PVX), whose virions have been detected in the apoplast but not associated with EVs (11), this is, to our knowledge, the first report of infectious viral entities isolated from the apoplast in association with EVs. Furthermore, although deletion of CP does not abolish the accumulation of viral DNA in the EV-enriched P40 fraction—indicating that vesicle-associated export, as well as plant infection, can occur independently of complete virion formation—the requirement of CP for geminiviral infectivity suggests that viral DNA must be at least coated or stabilized by CP to remain competent for infection.

Although geminiviruses are traditionally thought to rely exclusively on plasmodesmata for cell-to-cell movement, our results raise the possibility that, in addition to this symplastic route, they may exploit other pathways for genome dissemination, including extracellular routes. In this scenario, EVs could mediate short-range movement through the apoplast. The existence of redundant viral transport systems would increase the robustness of the viral life cycle and may contribute to the high efficiency of plant infection. Furthermore, an apoplastic movement pathway would provide an additional mechanism to infect cells that are not directly connected by plasmodesmata or in which plasmodesmal gating cannot be efficiently manipulated by the virus.

To participate in viral intercellular transmission, EVs must be capable of crossing the plant cell wall. Although the plant cell wall has traditionally been viewed as a barrier to such trafficking, there is evidence suggesting that it can become permeable under certain conditions. Thus, using focused high-resolution scanning electron microscopy (FIB-EHRSEM), Movahed et al. 2019 (1) showed that in TuMV-infected plants, extracellular vesicles move within the cell wall, indicating that it does not represent an absolute obstacle. Since the size exclusion limit of the plant cell wall diffusion has been estimated to be in the nanometer range below the sizes of the EVs (∼30–150 nm or larger) (14), their mobility cannot be accounted for by simple diffusion, but rather implies local wall remodelling/deformation, or the existence of larger, transient pores. Plant cell wall permeability is largely determined by the composition and organization of its polysaccharide matrix, particularly the degree of homogalacturonan methylesterification, which modulates cell wall stiffness, porosity, and its mechanical properties. This process is controlled by the opposing activities of pectin methylesterases (PMEs) and their inhibitors (PMEIs), which together control cell wall plasticity (53, 54). Accumulating evidence identifies PMEs as key host factors during viral infection (55): (i) their expression is induced by multiple viruses, including TuMV, cucumber mosaic virus, tobacco mosaic virus (TMV), and the geminivirus CabLCV (56–59), (ii) the interaction between the movement protein of TMV and the cell wall-associated PME contributes to local and systemic viral movement (56, 60, 61) and (iii) reduction of PME activity, either by silencing or PMEI overexpression, restricts viral spread of TMV and turnip vein clearing virus (62, 63). Given that the organization and density of the pectin network can vary spatially and dynamically, creating regions of differential permeability in the cell wall, PME induction during viral infection could remodel the wall structure, increasing its porosity and potentially facilitating the passage of large structures such as extracellular vesicles. Consistent with this, multiple cell wall-remodelling enzymes were detected in the CabLCV-infected P40 proteome (PMEs, PMEIs, expansins, pectin acetylesterases, pectin lyases, glucan-endo-glucosidases, etc.) supporting the notion that the plant cell wall architecture is actively modulated through apoplastic secretion (Dataset S1).

### Why is viral EV loading conserved in plants?

The growing evidence for viral secretion into the plant apoplast during both DNA and RNA virus infections suggests that this process is conserved and functionally relevant to the viral life cycle. Beyond its roles within the plant, the significance of apoplastic release and EV association becomes particularly compelling when considered in the context of geminiviruses as arboviruses (arthropod-borne viruses), an informal term referring to viruses transmitted by arthropod vectors.

The life cycle of plant circulative-transmitted arboviruses (viruses acquired by arthropod vectors that move internally through the vector before transmission to a new plant) requires adaptation to two fundamentally distinct cellular environments: plant hosts and arthropod vectors. In plants, viruses move within a rigid, cell wall–constrained system, relying predominantly on symplastic transport through plasmodesmata, whose permeability is actively modulated by viral and host factors. In contrast, arthropod vector cells lack cell walls and cytoplasmic continuity, forcing viruses to navigate extracellular spaces via endocytic and exocytic pathways. During circulative transmission in arthropod vectors, viruses must cross multiple physiological barriers, including the midgut, hemolymph, salivary glands, and ovary, through coordinated cycles of entry, trafficking, and release. Consistent with this, clathrin-mediated endocytosis facilitates begomovirus entry into vector midgut cells (64, 65), with early endosomes supporting subsequent intracellular trafficking (66). Similar mechanisms operate in other geminiviruses, such as members of the genus *Mastrevirus*, where capsid protein has been detected within membranous vesicles in vector midgut cells (67). Vesicle-mediated viral egress has been also described for plant reoviruses, such as rice dwarf virus, which exploits MVB–derived exosomes to reach salivary cavities prior to transmission into the plant phloem (68).

Collectively, these observations support a model in which successful arboviral infection relies on a dual transport strategy: symplastic movement within plants and vesicle-mediated trafficking adapted to extracellular environments in arthropod vectors. From this perspective, EV-mediated viral export in plants may not be incidental but instead reflect an evolutionarily conserved trait selected to optimize transmission across kingdoms. Notably, this raises the possibility that apoplastic viral accumulation in plants represents a transmission-adapted process, with intracellular spread and extracellular release forming complementary components of the viral life cycle. Whether EV association confers a direct advantage within plants, enhances vector acquisition, or instead acts as a regulatory or decoy mechanism remains unresolved. In this regard, it will be important to determine whether vesicle-associated viral components promote uptake by vectors or, alternatively, interfere with transmission, analogous to the role of bacterial outer membrane vesicles in modulating phage infection (69).

Taken together, our results support a revised framework for begomovirus biology as insect-transmitted viruses, in which symplastic and apoplastic pathways are not mutually exclusive but functionally integrated components of the viral life cycle. By coupling intracellular movement through plasmodesmata with EV-mediated trafficking across extracellular spaces, geminiviruses may achieve the flexibility required to propagate within plant tissues while remaining pre-adapted for transmission through arthropod vectors. This dual transport strategy provides a mechanistic bridge between host infection and vector-mediated spread, and positions extracellular vesicles as central players at the plant–vector interface. Elucidating how these pathways are coordinated will be critical to understand viral fitness, transmission efficiency, and the evolution of cross-kingdom infection strategies.

## Materials and Methods

### Microorganisms. Bacterial strains and infectious clones

Standard methods in microbiology were followed for bacterial manipulation (70, 71). *Agrobacterium tumefaciens* strain GV3101(pMP90) was employed for the delivery of the geminiviral infectious clones of cabbage leaf curl virus (CabLCV, GenBank accessions U65529 for DNA A and U65530 for DNA B) and beet curly top virus (BCTV, GenBank accession NC001412).

### Plant material and growth conditions

The *Arabidopsis thaliana* (L.) Heynh genotypes used were Columbia-0 (Col-0), both wild-type and 35S::RFP-PEN1×35S::TET8-GFP (9). Arabidopsis plants were grown in controlled growth chambers at 22 ºC, 60 % relative humidity and short-day conditions (150 μmol/m^2^s light for 8 h per day). Seeds were sterilized using 10 % v/v bleach (30 s), then ethanol 70 % v/v (2 min) and washed three times with sterile deionized water. Seeds were subsequently stratified for at least 2 days at 4 ºC in the dark and germinated on MS media 1 % agar pH 5.8 (Duchefa Biochemie M0231.0050) in the same growth conditions.

### Geminivirus agroinoculation assays

For Arabidopsis agroinoculation, 5-6-week-old Arabidopsis plants were injected with the infectious clones of CabLCV (72) or BCTV (73) as described in Ascencio-Ibáñez et al., 2008 (59). For *Nicotiana benthamiana* Domin agroinoculation, 5-week-old plants were injected with infectious clones of CabLCV (72). *Agrobacterium tumefaciens* clones harboring the corresponding infectious clones were grown overnight on LB liquid medium supplemented with the proper antibiotics. Saturated bacterial cultures were pelleted at 4,000 × g for 10 min and resuspended in agroinoculation buffer (10 mM MgCl2, 10 mM MES pH 5.6, and 100 μM acetosyringone), adjusting optical density to 1 (OD_600_ = 1) and incubating in the dark for 2 h to 4 h.

Arabidopsis apical leaves were punctured using insulin syringes, poking 20-30 holes per plant, and a droplet of *A. tumefaciens* suspension was deposited over the punctate surface. Inoculated plants were covered with plastic domes for 2 days. For *N. benthamiana* inoculation, bacterial cultures were inoculated using insulin syringes in the axillary bud of the fourth/fifth youngest leaf of the plant.

### Geminivirus mechanical inoculation assays

Arabidopsis mechanical inoculation was achieved through carborundum abrasion. Source inoculum was extracted from symptomatic young leaves from CabLCV-infected plants at 21 days post-infection. Infected plant tissue was macerated in 0.01 M phosphate buffer solution pH 7.0 to a final concentration of 50 mg/mL and centrifuged at 16 000 × g for 10 min at 4 ºC. Supernatant was transferred to new tubes and kept on ice.

4-5-week-old Arabidopsis rosettes were sprinkled over with carborundum powder and 150 μL of inoculum was rubbed over the surface of every leaf on the rosette using gloves. Inoculated plants were covered under a plastic dome from 24 h to 48 h. For P40 infectivity assays, at least the equivalent of 20 mL of AWF isolated from CabLCV-infected plants was inoculated.

### Isolation of apoplastic wash fluid (AWF)

Apoplastic wash fluid (AWF) from Arabidopsis was isolated following a previously published protocol (18). Briefly, 6–8-week-old Arabidopsis rosettes were collected from the soil after root excision and rinsed in distilled water three times. Rosettes were subsequently vacuum infiltrated (SEGOVAC 6.0) in a beaker once submerged abaxial side up in vesicle isolation buffer (VIB, 20 mM MES, 2 mM CaCl2 and 10 mM NaCl pH 6). Vacuum was applied for 1 min until most of the leaf area was clearly infiltrated. Infiltrated rosettes were gently patted dry using paper towels to remove the excess buffer. Rosettes were then introduced adaxial side up in modified 50 mL syringes with 10 extra holes in the bottom to allow free buffer flowthrough. The loaded syringes were placed in 250 mL bottles for centrifugation in a fixed-angle rotor (HIMAC R16A3) at 700 × g for 30 min at 4 ºC using slow acceleration to isolate AWF from the leaves. AWF was passed through a 0.45 μm filter.

For *N. benthamiana* AWF isolation, we followed a similar methodology including some modifications. Fully expanded leaves were harvested, rinsed three times with sterile deionized water and vacuum infiltrated with VIB as described above. *N. benthamiana* leaves were dry patted, rolled and introduced stem cut site facing up into modified 50 mL syringes for centrifugation and filtering in the same conditions as previously detailed.

### EV pellet isolation

EV isolation from AWF was performed as previously published in Rutter et al. 2017 (18). Filtered AWF was centrifuged (Eppendorf F45-24-11) at 10 000 × g for 30 min at 4 ºC, then the supernatant was transferred to new tubes and centrifuged at 40 000 × g for 1 h at 4 ºC. The resulting supernatant was discarded by decanting, and the P40 pellet was resuspended in 20 mM Tris HCl pH 7.5.

### Nanoparticle tracking analysis (NTA)

Resuspended EV pellets were immediately used for NTA after isolation (without freezing). Nanoparticle tracking analysis (NTA) was conducted using a nanoparticle tracker (Particle Metrix PMX 120 Zetaview Mono 488 Laser running Zetaview software version 8.05.16 SP) as previously described in Koch et al., 2025 (9).

### Confocal laser scanning microscopy (CLSM)

Resuspended EV pellets from transgenic reporter lines RFP-PEN1 × TET8-GFP were visualized under CLSM after EV isolation (without freezing). EV samples fluorescence was detected using Zeiss LSM880 confocal microscope (Zeiss, http://www.zeiss.com). GFP and RFP fluorescence were observed by 488 nm with an argon laser and 561 nm excitation, respectively. Fluorophore emission was examined with band-pass filters BP525/50 (495 nm to 525 nm) and BP605/70 (582 nm to 651 nm), respectively.

### Transmission electron microscopy (TEM)

Resuspended EV pellets were visualized under TEM after EV isolation (without freezing) as previously published in Koch et al., 2025 (9).

### Protein preparation and immunoblotting

Plant tissue preparation for protein extraction and immunoblotting was performed as described in Koch et al., 2025 (9). In brief, leaves were snap-frozen in liquid nitrogen and macerated using plastic pestles until they were ground into a fine powder. Cold protein extraction buffer (150 mM NaCl, 50 mM Tris HCl pH 7.5, 0.1% v/v Nonidet-P40, 1% v/v plant protease inhibitor cocktail) was added to an equivalent mass of macerated tissue and the mix was centrifuged at 16,000 × g for 10 min at 4 ºC. Supernatant was transferred to new tubes for protein quantification using Pierce 660 nm reagent (Sigma #22660) according to the manufacturer’s instructions and comparing to a bovine serum albumin calibration curve.

For immunoblotting, samples were mixed with 5X loading dye (250 mM Tris-HCl pH 6.8, 10% w/v SDS, 40% v/v glycerol, 0.02% bromoethanol blue, 5% v/v 2-mercaptoethanol) and incubated at 95 ºC for 5 min. Protein samples were then separated on 10 % SDS-PAGE gels. Proteins were then transferred to 0.45 μm PVDF and visualized by applying Ponceau Stain (0.1% w/v Ponceau S Stain, 5% v/v acetic acid) for 3 min and rinsing in distilled water or Coomassie Brilliant Blue (0.1 % w/v Coomassie R-250, 40 % v/v ethanol, 10 % v/v acetic acid) for 10 minutes and rinsing three times with destaining solution (40 % v/v ethanol, 10 % v/v acetic acid). Protein immunodetection was performed according to standard protocols and using the antibodies indicated in Table S4. Membranes were developed after incubation with enhanced chemiluminescent substrate (National Diagnostics Protoglow #CL-300). Densitometry comparative analyses were processed using FIJI.

### Nucleic acid isolation

Plant cell lysate DNA was extracted from Arabidopsis leaves following the CTAB method as described in Lukowitz et al., 1996 (74). In brief, approximately 50-100 mg of plant ground tissue was homogenized into 400 μL of extraction buffer (2 % cetyl trimethylammonium bromide (CTAB), 1.5 M NaCl, 100 mM Tris pH 8, 100 mM EDTA pH 8) and incubated at 65 ºC for 15 min. After cooling, the mixture was extracted with chloroform/isoamyl alcohol (24:1) and the nucleic acids were precipitated with isopropanol. DNA was finally resuspended in water and treated with RNase (10 mg/mL) (Invitrogen).

### Polymerase chain reaction (PCR) and Quantitative real-time PCR (qPCR)

Polymerase chain reaction (PCR) amplification was performed following standard procedures and using specific primers included in Table S5. Quantitative real-time PCR was employed for viral DNA quantification. Template DNA containing 1-10 ng was mixed in 10 μL reactions including specific primers at 10 μM (Table S5), and 5 μL of SsoFast EvaGreen Supermix (Bio-Rad). The reactions were carried out in CFX96 and CFX384 Touch Real-Time PCR Detection System instruments (Bio-Rad) following 10 min at 95 ºC and 40 cycles of 15 s at 95 ºC and 10 s at 60 ºC. Three technical replicates per sample were included in each reaction plate. The data was analysed using Livak’s 2-DDCT method (75).

### Rolling circle amplification (RCA)

Complete geminiviral genomes were amplified using the phi29-XT RCA kit (New England Biolabs, #E1603) following the manufacturer’s protocol. Briefly, DNA templates were mixed in 18 μL reactions containing exonuclease-resistant random primers and incubated at 100 ºC for 3 minutes. Then, 2 μL of phi29-XT DNA Polymerase was added to the reaction and the whole volume was incubated at 42 ºC for 2 hours, followed by 10 minutes at 65 ºC for DNA polymerase inactivation. RCA products were analysed by gel electrophoresis upon specific endonuclease restriction treatment.

### DNase and trypsin protection assays

For DNase and protease protection assays, EV pellets were respectively resuspended in 20 mM and 150 mM Tris HCl pH 7.8 buffer and immediately aliquoted after P40 isolation. For sonication treatment, samples were immersed in a cold bath sonicator (J.P. Selecta, #3000512), exposed to ultrasounds for 15 minutes and then heat-shocked at 95 ºC for 5 min. For DNase treatment, Recombinant DNase I (RNase-free) (Takara, #2270A) was added to samples to a final concentration of 0.25 units/μL and incubated at 37 ºC for 45 min. DNase reaction stopped after the addition of 0.5 μL of proteinase K (ThermoFisher Scientific #EO0491) and further incubation at 37 ºC for 10 min followed by a final denaturation step at 80 ºC for 5 min. For protease treatment, samples were treated with trypsin (Promega #V5113) to a final concentration of 10–20 μg/mL, and samples were incubated for 1 h at 37 ºC.

### Mass spectrometry (MS)

P40 samples from CabLCV-infected and non-infected Arabidopsis plants were collected in three independent biological replicates. For protein digestion, each sample was diluted with lysis buffer to 2.5% sodium dodecyl sulfate (SDS) and 25 mM triethylammonium bicarbonate (TEAB). Samples were reduced and alkylated by adding 5 mM tris(2-carboxyethyl)phosphine (TCEP) and 10 mM chloroacetamide (CAA) for 30 minutes at 60°C. The protein extracts were centrifuged at 18,000 × g for 10 min and the supernatant was transferred to a new tube and quantified using PIERCE 660 nm reagent (compatible with ionic detergents). Protein digestion on S-Trap columns (Protifi) was performed following the manufacturer’s instructions with minor changes (76). Each sample was digested at 37°C overnight using trypsin:protein ratio of 1:15. Tryptic peptides were clean-up using a StageTip C18 prior to LC-ESI-MS/MS analysis (77). Eluted peptides were dried in a speed vacuum and quantified by fluorimetry (QuBit) according to the manufacturer’s instructions. For nano-Liquid Chromatography coupled to Electrospray Ionization Tandem Mass Spectrometry (nanoLC-ESI-MS/MS) analysis, 500 ng of each sample was individually analyzed using an Ultimate 3000 nano HPLC system (Thermo Fisher Scientific) coupled online to an Orbitrap Exploris™ 240 mass spectrometer (Thermo Fisher Scientific). Each sample (500 ng in 5 µl of sample resuspended in mobile phase A: 0.1% formic acid (FA)) was loaded on a 50 cm × 75 μm Easy-spray PepMap C18 analytical column at 45°C and separated at a flow rate of 250 nL/min using a 120 min gradient ranging from 2 % to 95 % mobile phase B (80 % acetonitrile (ACN) in 0.1% FA). Data acquisition was performed using a data-dependent top-20 method, in full scan positive mode, scanning 375 to 1200 m/z. MS1 scans were acquired at an orbitrap resolution of 60,000 at m/z 200, with normalized automatic gain control (AGC) target of 300%, a radio frequency (RF) lens of 80%, and an automatic maximum injection time (IT). The 20 most intense ions from each MS1 scan were selected and fragmented with a higher-energy collisional dissociation (HCD) of 30%. Resolution for HCD spectra was set to 15,000 at m/z 200, with a normalized AGC target of 50% and an automatic maximum IT. Isolation of precursors was performed with an isolation window of 1 m/z and 10 s of exclusion duration. Precursor ions with charge states from 2+ to 5+ were included. Raw data files were processed using the Proteome Discoverer 2.5.0.400 software (Thermo Scientific, Bremen, Germany), and a database search was carried out using using three search engines (Mascot (v2.8.1), MsFragger (v3.8), and Sequest HT) against a Arabidopsis thaliana UniProtKB reviewed database (20250424, 39,366 sequences) containing the most common laboratory contaminants (cRAP database with 70 sequences). Search parameters were set as follows: cysteine carbamidomethylation (+57.021464 Da); methionine oxidation (M) (+15.994915 Da), N-term acetylation (+42.010565 Da), and Gln→pyro-Glu (-17.026549 Da) as variable modifications. Precursor mass tolerances were set at 10 ppm and the fragment mass tolerance at 0.02 Da and trypsin/P was selected as a protease with a maximum of 2 missed cleavage sites. False discovery rate (FDR) was calculated using the processing node Percolator (maximum delta Cn 0.05; decoy database search target) and the validation of proteins, peptides, and peptide spectral matches (PSMs) peptides with an FDR≤1 %. Precursor ion quantitation was also performed in Proteome Discoverer using the “Minora” feature in the processing method and the “Feature Mapper” and “Precursor Ions Quantifier” nodes in the consensus step. The Feature Mapper node in the consensus method was used to create features from unique peptide-specific peaks by applying a chromatographic retention time alignment with a maximum shift of 10 min and a threshold of signal-to-noise of 5. The following parameters were set in the “Precursor Ions Quantifier” node: unique+razor peptides were used for quantitation, precursor abundance was based on intensity, and normalization mode was based on the total peptide amount. Protein abundances were calculated by summing sample abundances of the connected peptide groups. Protein abundances were calculated by summing sample abundances of the connected peptide groups. Pairwise-based ratios were calculated and a t-test background-based was performed as a statistical test. To determine the differentially expressed proteins in each comparison, the following filters were considered: quantitation by protein group (only master proteins were selected) with a protein FDR confidence of high, normalized protein abundance was detected in at least 2 samples, the percentage of coefficient of variation of abundance values should have a value in each sample groups, and the p-value adjusted using Benjamini–Hochberg correction was set ≤0.05.

### Image analysis of fluorescent foci

To quantify RFP-PEN1 and TET8-GFP foci in P40 pellets from from non-infected and CabLCV-infected plants, images were analyzed using the FIJI software (https://fiji.sc/) (78). Binary masks were generated by identifying local maxima by applying the *FindMaxima* command. The resulting foci counts were relativized to the respective non-infected treatment within each biological replicate.

### Statistical analysis

Statistically significant differences between the number of nanoparticles per mL of AWF detected by NTA and fluorescent RFP-PEN1 and TET8-GFP foci in P40 pellets from non-infected and CabLCV-infected plants were evaluated using a ratio paired t-test (****p-value < 0.0001; *p-value < 0.05).

## Acknowledgments

This work was supported by the Spanish Ministerio de Economía y Competitividad, co-financed by the European Regional Development Fund, ERDF (grant PID2022-139376OB-C31 to A.G.C. and E.R.B.) and Spanish Ministerio de Ciencia, Innovación y Universidades (FPU fellowship number FPU20/03715 to P.M.M.). This work was also supported by grants from the U.S. National Science Foundation Plant Biotic Interactions and Plant Genome Research programs (grant numbers IOS-1645745, IOS-1842685, and IOS-2141969 to R.W.I.) and by the Novo Nordisk Foundation. B.L.K. was supported by a Carlos O. Miller Graduate Fellowship, the Colonel Bayard Franklin Floyd Memorial Fund, and the William R. Ogg Fellowship, all from the IU Foundation. We thank CNB Proteomics Facilities and M. Lucía Borniego and Meenu Singla-Rastogi for advice on EV isolation and characterization

