## Supporting information for "Extracellular vesicles associate with infectious geminiviral particles in the apoplast of infected plants"

* Eduardo R. Bejarano

*Araceli G. Castillo

**Figures**

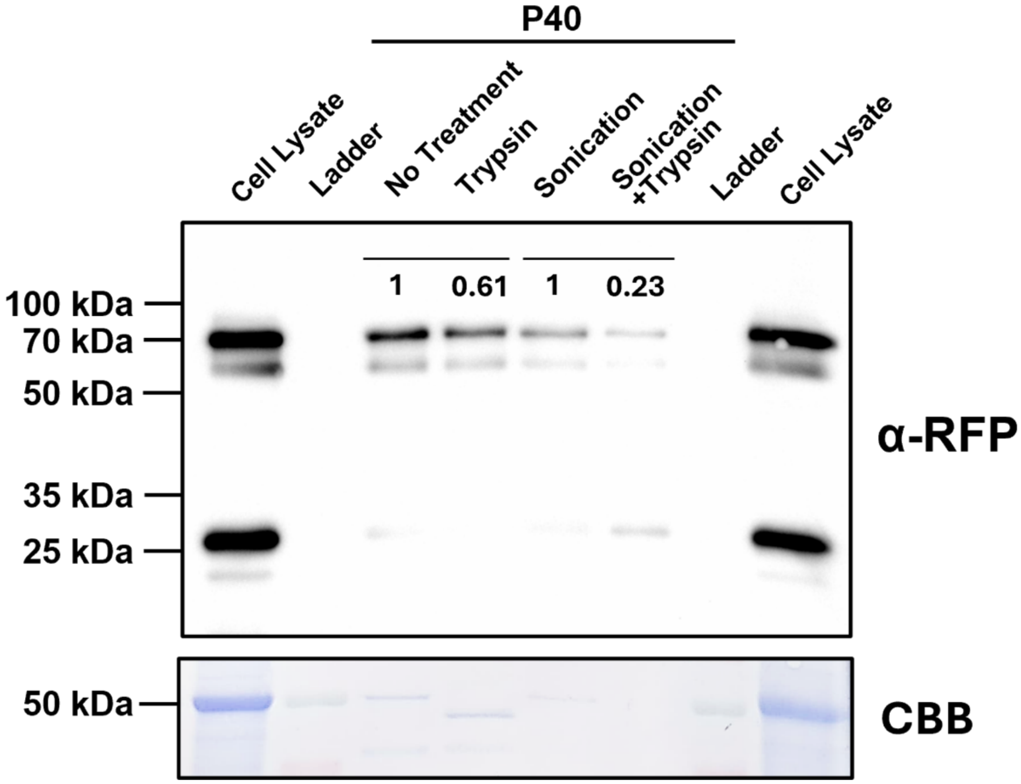


**Fig. S1**. Sonication treatment disrupts EV structure and enhances trypsin-mediated RFP-PEN1 proteolysis. P40 pellets subjected to trypsin incubation (10-20 µg/mL) for 1 h at 37 ºC after sonication treatment showed a 2–3-fold reduction in RFP-PEN signal compared to non-sonicated P40 pellets, suggesting an effect of ultrasounds in disrupting EVs. Experiments were reproduced twice with similar results.

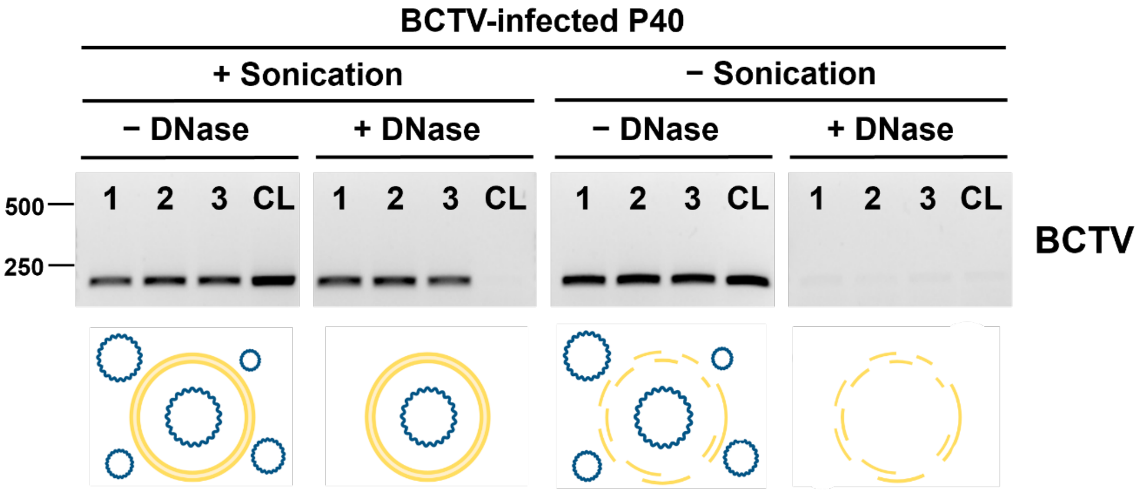


**Fig. S2.** Protection of geminiviral DNA sequences in P40 fractions is also observed for geminiviruses belonging to different species and genera. Geminiviral sequences in P40 fractions were protected against DNase treatment in *Arabidopsis thaliana* plants infected with BCTV (genus *Curtovirus*). **1-3**: independent P40 fractions from the same batch of AWF. **CL**: Cell lysate DNA extraction from BCTV-infected plants. Experiments were reproduced twice with similar results.

### DNA A alignment

```
DNA_A-CabLCV[U65529]      ttttgtaaataatatttaggactccaggggaggacgccaggagagctctctctaaaacct      60
RCA_P40-CabLCV-Clone1     ttttgtaaataatatttaggactccaggggaggacgccaggagagctctctctaaaacct      60
                          ************************************************************

DNA_A-CabLCV[U65529]      attgtttttggtgtcttggtgtccaatatatactaaaagcctctggggaacaccaggggc      120
RCA_P40-CabLCV-Clone1     attgtttttggtgtcttggtgtccaatatatactaaaagcctctggggaacaccaggggc      120
                          ************************************************************

DNA_A-CabLCV[U65529]      aaaagcggccatccgcaataatattaccggatggccgcgatttttttggtgtcttgcgtg      180
RCA_P40-CabLCV-Clone1     aaaagcggccatccgcaataatattaccggatggccgcgatttttttggtgtcttgcgtg      180
                          ************************************************************

DNA_A-CabLCV[U65529]      ggaccacgcattaaatgaaatctaaccaatcacatgccgcctgacaaggttagctattaa      240
RCA_P40-CabLCV-Clone1     ggaccacgcattaaatgaaatctaaccaatcacatgccgcctgacaaggttagctattaa      240
                          ************************************************************

DNA_A-CabLCV[U65529]      ggctaactgagtgcgctgtgggcctatataaagacatgtgatttcaatttatgctttact      300
RCA_P40-CabLCV-Clone1     ggctaactgagtgcgctgtgggcctatataaagacatgtgatttcaatttatgctttact      300
                          ************************************************************

DNA_A-CabLCV[U65529]      tcgaaatgcctaagcgggatgccccgtggcgttctatggcggggacctctaaagtgtccc      360
RCA_P40-CabLCV-Clone1     tcgaaatgcctaagcgggatgccccgtggcgttctatggcggggacctctaaagtgtccc      360
                          ************************************************************

DNA_A-CabLCV[U65529]      gcaatgctaactattcacctcgtgcaggtatgatccataaatttgataaggccgctgctt      420
RCA_P40-CabLCV-Clone1     gcaatgctaactattcacctcgtgcaggtatgatccataaatttgataaggccgctgctt      420
                          ************************************************************

DNA_A-CabLCV[U65529]      gggttaacaggcccatgtacaggaagcccaggatttataggacgtttagaagcccagatg      480
RCA_P40-CabLCV-Clone1     gggttaacaggcccatgtacaggaagcccaggatttataggacgtttagaagcccagatg      480
                          ************************************************************

DNA_A-CabLCV[U65529]      ttcctagaggctgtgaagggccttgcaaagtccagtcttatgagcagcggcatgacattt      540
RCA_P40-CabLCV-Clone1     ttcctagaggctgtgaagggccttgcaaagtccagtcttatgagcagcggcatgacattt      540
                          ************************************************************

DNA_A-CabLCV[U65529]      cccatgttggtaaggtcatgtgtatttcagacataacacgtggtaatggtattacccatc      600
RCA_P40-CabLCV-Clone1     cccatgttggtaaggtcatgtgtatttcagacataacacgtggtaatggtattacccatc      600
                          ************************************************************

DNA_A-CabLCV[U65529]      gtgttgggaagcgtttttgcgtcaagtccgtgtacattttgggcaagatatggatggacg      660
RCA_P40-CabLCV-Clone1     gtgttgggaagcgtttttgcgtcaagtccgtgtacattttgggcaagatatggatggacg      660
                          ************************************************************

DNA_A-CabLCV[U65529]      agaatatcaagctcaagaaccacaccaacagtgtcatgttttggttggttagggacagaa      720
RCA_P40-CabLCV-Clone1     agaatatcaagctcaagaaccacaccaacagtgtcatgttttggttggttagggacagaa      720
                          ************************************************************

DNA_A-CabLCV[U65529]      gaccatatggcacccctatggagtttggccaagtgttcaacatgttcgacaacgagccca      780
RCA_P40-CabLCV-Clone1     gaccatatggcacccctatggagtttggccaagtgttcaacatgttcgacaacgagccca      780
                          ************************************************************

DNA_A-CabLCV[U65529]      gtactgctactgtgaagaacgatcttcgtgatcgttatcaagtcatgcacaagttttacg      840
RCA_P40-CabLCV-Clone1     gtactgctactgtgaagaacgatcttcgtgatcgttatcaagtcatgcacaagttttacg      840
                          ************************************************************

DNA_A-CabLCV[U65529]      caaaggtaaccggtgggcagtatgcgagtaacgagcaggcgttggtgaagcgtttttgga      900
RCA_P40-CabLCV-Clone1     caaaggtaaccggtgggcagtatgcgagtaacgagcaggcgttggtgaagcgtttttgga      900
                          ************************************************************

DNA_A-CabLCV[U65529]      aggtcaataactacgttgtgtacaaccatcaagaagcagggaaatacgagaatcatacgg      960
RCA_P40-CabLCV-Clone1     aggtcaataactacgttgtgtacaaccatcaagaagcagggaaatacgagaatcatacgg      960
                          ************************************************************

DNA_A-CabLCV[U65529]      agaacgctctgttattgtatatggcatgtactcatgcctctaatcctgtgtatgcgacat      1020
RCA_P40-CabLCV-Clone1     agaacgctctgttattgtatatggcatgtactcatgcctctaatcctgtgtatgcgacat      1020
                          ************************************************************

DNA_A-CabLCV[U65529]      taaaaattcggatctatttttacgattcgataacaaattaataaattttgaaatttatta      1080
RCA_P40-CabLCV-Clone1     taaaaattcggatctatttttacgattcgataacaaattaataaattttgaaatttatta      1080
                          ************************************************************

DNA_A-CabLCV[U65529]      catgattctcgtgaacatgagttacataagatctgtccgttgcgaaacgaacagctctaa      1140
RCA_P40-CabLCV-Clone1     catgattctcgtgaacatgagttacataagatctgtccgttgcgaaacgaacagctctaa      1140
                          ************************************************************

DNA_A-CabLCV[U65529]      ttacatgattaataccaataacacctaggttatctaaatgagacattacaaggcatttga      1200
RCA_P40-CabLCV-Clone1     ttacatgattaataccaataacacctaggttatctaaatgagacattacaaggcatttga      1200
                          ************************************************************
```

```
DNA_A-CabLCV[U65529]      atctacttaaatatgtcggcccagaagctctcatcgaagtcgtccagacttggaagttga      1260
RCA_P40-CabLCV-Clone1     atctacttaaatatgtcggcccagaagctctcatcgaagtcgtccagacttggaagttga      1260
                          ************************************************************

DNA_A-CabLCV[U65529]      agtaggctttgcggagatccaatgctttccgtaggttgtggttgaaccggacttggatgt      1320
RCA_P40-CabLCV-Clone1     agtaggctttgcggagatccaatgctttccgtaggttgtggttgaaccggacttggatgt      1320
                          ************************************************************

DNA_A-CabLCV[U65529]      ggtagatcctggttctggtgtacaacgggtcctctacgcgttgtatcctgaaatataggg      1380
RCA_P40-CabLCV-Clone1     ggtagatcctggttctggtgtacaacgggtcctctacgcgttgtatcctgaaatataggg      1380
                          ************************************************************

DNA_A-CabLCV[U65529]      gatttggaacctcccagataaaaacggaattctctgcctgagctacagtgatgctctccc      1440
RCA_P40-CabLCV-Clone1     gatttggaacctcccagataaaaacggaattctctgcctgagctacagtgatgctctccc      1440
                          ************************************************************

DNA_A-CabLCV[U65529]      cggtgcgtgaatccattatctgcgcaattgatgtggaggaagatagaacacccgcagttc      1500
RCA_P40-CabLCV-Clone1     cggtgcgtgaatccattatctgcgcaattgatgtggaggaagatagaacacccgcagttc      1500
                          ************************************************************

DNA_A-CabLCV[U65529]      aaatcaatgcgtcttcggcgtacagctctccttttagcaatcttgtgctgtgctttgata      1560
RCA_P40-CabLCV-Clone1     aaatcaatgcgtcttcggcgtacagctctccttttagcaatcttgtgctgtgctttgata      1560
                          ************************************************************

DNA_A-CabLCV[U65529]      gaggggggcttcaagagtgatgaattttgcatttttggtagtccacgctcttagtgatgc      1620
RCA_P40-CabLCV-Clone1     gaggggggcttcaagagtgatgaattttgcatttttggtagtccacgctcttagtgatgc      1620
                          ************************************************************

DNA_A-CabLCV[U65529]      attttcctctttgttgaggaaacttatataactgctcccctctcctggattgcacagcac      1680
RCA_P40-CabLCV-Clone1     attttcctctttgttgaggaaacttatataactgctcccctctcctggattgcacagcac      1680
                          ************************************************************

DNA_A-CabLCV[U65529]      gattgagggtatgccacctttaatttgaactggcttgccgtacttacagtttgattgcca      1740
RCA_P40-CabLCV-Clone1     gattgagggtatgccacctttaatttgaactggcttgccgtacttacagtttgattgcca      1740
                          ************************************************************

DNA_A-CabLCV[U65529]      gtccctttgggccccaataagctctttccagtgctttagctttagataatgcggagctat      1800
RCA_P40-CabLCV-Clone1     gtccctttgggccccaataagctctttccagtgctttagctttagataatgcggagctat      1800
                          ************************************************************

DNA_A-CabLCV[U65529]      gtcatcaatgacgttatactccgcattatttgaaaagacctttgaattaaagtcgaggtg      1860
RCA_P40-CabLCV-Clone1     gtcatcaatgacgttatactccgcattatttgaaaagacctttgaattaaagtcgaggtg      1860
                          ************************************************************

DNA_A-CabLCV[U65529]      cccactcaaataattatgtggtcctaaagcacgcgcccacatggtcttgccggttcgtga      1920
RCA_P40-CabLCV-Clone1     cccactcaaataattatgtggtcctaaagcacgcgcccacatggtcttgccggttcgtga      1920
                          ************************************************************

DNA_A-CabLCV[U65529]      atcaccttcaactatgatactaataggtctttccgcccgcgcagcggcactccgaccaaa      1980
RCA_P40-CabLCV-Clone1     atcaccttcaactatgatactaataggtctttccgcccgcgcagcggcactccgaccaaa      1980
                          ************************************************************

DNA_A-CabLCV[U65529]      atagtcatctgcccatgaactcatctcgtccgggacgttagtaaaggaggagagttgaaa      2040
RCA_P40-CabLCV-Clone1     atagtcatctgcccatgaactcatctcgtccgggacgttagtaaaggaggagagttgaaa      2040
                          ************************************************************

DNA_A-CabLCV[U65529]      cggaggagcccatggttccggagccttagtaaagagtcgctctatgttagctctaacatt      2100
RCA_P40-CabLCV-Clone1     cggaggagcccatggttccggagccttagtaaagagtcgctctatgttagctctaacatt      2100
                          ************************************************************

DNA_A-CabLCV[U65529]      atgataactaacaatgaacgtctttggatctccagcccttataattgcgagagcctcttc      2160
RCA_P40-CabLCV-Clone1     atgataactaacaatgaacgtctttggatctccagcccttataattgcgagagcctcttc      2160
                          ************************************************************

DNA_A-CabLCV[U65529]      cacacatcccgcattgacggcgttgtggtagacgtcgtctttatttgcctttgtaccccc      2220
RCA_P40-CabLCV-Clone1     cacacatcccgcattgacggcgttgtggtagacgtcgtctttatttgcctttgtaccccc      2220
                          ************************************************************

DNA_A-CabLCV[U65529]      agacaccttgtactgcccggattcacaataatcaccatctttggtgatgtaattcttgac      2280
RCA_P40-CabLCV-Clone1     agacaccttgtactgcccggattcacaataatcaccatctttggtgatgtaattcttgac      2280
                          ************************************************************

DNA_A-CabLCV[U65529]      ggcattggtgtctttggctgcctgaatgtttgggtgaaaattggcagaccttctggggtg      2340
RCA_P40-CabLCV-Clone1     ggcattggtgtctttggctgcctgaatgtttgggtgaaaattggcagaccttctggggtg      2340
                          ************************************************************

DNA_A-CabLCV[U65529]      agtgatgtcgaaaaatctagcatccttgatgttcgactttcctgatagttggatgagaca      2400
RCA_P40-CabLCV-Clone1     agtgatgtcgaaaaatctagcatccttgatgttcgactttcctgatagttggatgagaca      2400
                          ************************************************************
```

```
DNA_A-CabLCV[U65529]       gtgtaaatgggggaacccgtctgaatgttcctctcttgcgactctgatgtatgtgggttt    2460
RCA_P40-CabLCV-Clone1      gtgtaaatgggggaacccgtctgaatgttcctctcttgcgactctgatgtatgtgggttt    2460
                           ************************************************************

DNA_A-CabLCV[U65529]       gacgactgaccacgacagggtttgaagcatctgaagagcttcatctttgggtatgtcgca    2520
RCA_P40-CabLCV-Clone1      gacgactgaccacgacagggtttgaagcatctgaagagcttcatctttgggtatgtcgca    2520
                           ************************************************************

DNA_A-CabLCV[U65529]       ctggggatatgttaagaatatatttcgggctgctaaacgaaacgatttagggtttcgtgg    2580
RCA_P40-CabLCV-Clone1      ctggggatatgttaagaatatatttcgggctgctaaacgaaacgatttagggtttcgtgg    2580
                           ************************************************************

DNA_A-CabLCV[U65529]       cat    2583
RCA_P40-CabLCV-Clone1      cat    2583
                           ***
```

### DNA B Alignment

```
DNA_B-CabLCV[U65530]       ttatagaaatccgaccgcgcagcggcaggggcttcaataatatctgggtttagtaaaaac    60
RCA_P40-CabLCV-Clone2      ttatagaaatccgaccgcgcagcggcaggggcttcaataatatctgggtttagtaaaaac    60
RCA_P40-CabLCV-Clone3      ttatagaaatccgaccgcgcagcggcaggggcttcaataatatctgggtttagtaaaaac    60
RCA_P40-CabLCV-Clone4      ttatagaaatccgaccgcgcagcggcaggggcttcaataatatctgggtttagtaaaaac    60
RCA_P40-CabLCV-Clone5      ttatagaaatccgaccgcgcagcggcaggggcttcaataatatctgggtttagtaaaaac    60
RCA_P40-CabLCV-Clone6      ttatagaaatccgaccgcgcagcggcaggggcttcaataatatctgggtttagtaaaaac    60
                           ************************************************************

DNA_B-CabLCV[U65530]       tgtgttgaagaacagaaaaatgaaacgacctgtttcgaaaggaggaagagagaaaatctg    120
RCA_P40-CabLCV-Clone2      tgtgttgaagaacagaaaaatgaaacgacctgtttcgaaaggaggaagagagaaaatctg    120
RCA_P40-CabLCV-Clone3      tgtgttgaagaacagaaaaatgaaacgacctgtttcgaaaggaggaagagagaaaatctg    120
RCA_P40-CabLCV-Clone4      tgtgttgaagaacagaaaaatgaaacgacctgtttcgaaaggaggaagagagaaaatctg    120
RCA_P40-CabLCV-Clone5      tgtgttgaagaacagaaaaatgaaacgacctgtttcgaaaggaggaagagagaaaatctg    120
RCA_P40-CabLCV-Clone6      tgtgttgaagaacagaaaaatgaaacgacctgtttcgaaaggaggaagagagaaaatctg    120
                           ************************************************************

DNA_B-CabLCV[U65530]       ggtttctaaacagaaagggaataacgatagttctaacgaaggatgagtatgttaggttat    180
RCA_P40-CabLCV-Clone2      ggtttctaaacagaaagggaataacgatagttctaacgaaggatgagtatgttaggttat    180
RCA_P40-CabLCV-Clone3      ggtttctaaacagaaagggaataacgatagttctaacgaaggatgagtatgttaggttat    180
RCA_P40-CabLCV-Clone4      ggtttctaaacagaaagggaataacgatagttctaacgaaggatgagtatgttaggttat    180
RCA_P40-CabLCV-Clone5      ggtttctaaacagaaagggaataacgatagttctaacgaaggatgagtatgttaggttat    180
RCA_P40-CabLCV-Clone6      ggtttctaaacagaaagggaataacgatagttctaacgaaggatgagtatgttaggttat    180
                           ************************************************************

DNA_B-CabLCV[U65530]       tcagataacatgagtataataatctaagactaagtctgattatatagagagcaaatttga    240
RCA_P40-CabLCV-Clone2      tcagataacatgagtataataatctaagactaagtctgattatatagagagcaaatttga    240
RCA_P40-CabLCV-Clone3      tcagataacatgagtataataatctaagactaagtctgattatatagagagcaaatttga    240
RCA_P40-CabLCV-Clone4      tcagataacatgagtataataatctaagactaagtctgattatatagagagcaaatttga    240
RCA_P40-CabLCV-Clone5      tcagataacatgagtataataatctaagactaagtctgattatatagagagcaaatttga    240
RCA_P40-CabLCV-Clone6      tcagataacatgagtataataatctaagactaagtctgattatatagagagcaaatttga    240
                           ************************************************************

DNA_B-CabLCV[U65530]       agaaattaaaaaagcttcgaatgttatgttagtggcatattcgtaaatatgttcatggac    300
RCA_P40-CabLCV-Clone2      agaaattaaaaaagcttcgaatgttatgttagtggcatattcgtaaatatgttcatggac    300
RCA_P40-CabLCV-Clone3      agaaattaaaaaagcttcgaatgttatgttagtggcatattcgtaaatatgttcatggac    300
RCA_P40-CabLCV-Clone4      agaaattaaaaaagcttcgaatgttatgttagtggcatattcgtaaatatgttcatggac    300
RCA_P40-CabLCV-Clone5      agaaattaaaaaagcttcgaatgttatgttagtggcatattcgtaaatatgttcatggac    300
RCA_P40-CabLCV-Clone6      agaaattaaaaaagcttcgaatgttatgttagtggcatattcgtaaatatgttcatggac    300
                           ************************************************************

DNA_B-CabLCV[U65530]       accaggagagctctcgtctaaaacctattgtttctggtgtcttggtgtcctataaatact    360
RCA_P40-CabLCV-Clone2      accaggagagctctcgtctaaaacctattgtttctggtgtcttggtgtcctataaatact    360
RCA_P40-CabLCV-Clone3      accaggagagctctcgtctaaaacctattgtttctggtgtcttggtgtcctataaatact    360
RCA_P40-CabLCV-Clone4      accaggagagctctcgtctaaaacctattgtttctggtgtcttggtgtcctataaatact    360
RCA_P40-CabLCV-Clone5      accaggagagctctcgtctaaaacctattgtttctggtgtcttggtgtcctataaatact    360
RCA_P40-CabLCV-Clone6      accaggagagctctcgtctaaaacctattgtttctggtgtcttggtgtcctataaatact    360
                           ************************************************************

DNA_B-CabLCV[U65530]       aaaagcctctggggactccataggacacgtgtacacatctcagcggccatccgtaataat    420
RCA_P40-CabLCV-Clone2      aaaagcctctggggactccataggacacgtgtacacatctcagcggccatccgtaataat    420
RCA_P40-CabLCV-Clone3      aaaagcctctggggactccataggacacgtgtacacatctcagcggccatccgtaataat    420
RCA_P40-CabLCV-Clone4      aaaagcctctggggactccataggacacgtgtacacatctcagcggccatccgtaataat    420
RCA_P40-CabLCV-Clone5      aaaagcctctggggactccataggacacgtgtacacatctcagcggccatccgtaataat    420
RCA_P40-CabLCV-Clone6      aaaagcctctggggactccataggacacgtgtacacatctcagcggccatccgtaataat    420
                           ************************************************************
```

```
DNA_B-CabLCV[U65530]       attaccggatggccgcaaatttttggagtcctggtgctggggaccatctcccgctcccct       480
RCA_P40-CabLCV-Clone2      attaccggatggccgcaaatttttggagtcctggtgctggggaccatctcccgctcccct       480
RCA_P40-CabLCV-Clone3      attaccggatggccgcaaatttttggagtcctggtgctggggaccatctcccgctcccct       480
RCA_P40-CabLCV-Clone4      attaccggatggccgcaaatttttggagtcctggtgctggggaccatctcccgctcccct       480
RCA_P40-CabLCV-Clone5      attaccggatggccgcaaatttttggagtcctggtgctggggaccatctcccgctcccct       480
RCA_P40-CabLCV-Clone6      attaccggatggccgcaaatttttggagtcctggtgctggggaccatctcccgctcccct       480
                           ************************************************************

DNA_B-CabLCV[U65530]       ttggcgcactctctctcatctctcgacgtggcgcgtcgaggagcgttggctaacgcttat       540
RCA_P40-CabLCV-Clone2      ttggcgcactctctctcatctctcgacgtggcgcgtcgaggagcgttggctaacgcttat       540
RCA_P40-CabLCV-Clone3      ttggcgcactctctctcatctctcgacgtggcgcgtcgaggagcgttggctaacgcttat       540
RCA_P40-CabLCV-Clone4      ttggcgcactctctctcatctctcgacgtggcgcgtcgaggagcgttggctaacgcttat       540
RCA_P40-CabLCV-Clone5      ttggcgcactctctctcatctctcgacgtggcgcgtcgaggagcgttggctaacgcttat       540
RCA_P40-CabLCV-Clone6      ttggcgcactctctctcatctctcgacgtggcgcgtcgaggagcgttggctaacgcttat       540
                           ************************************************************

DNA_B-CabLCV[U65530]       ctcggtgttaaacctttaatttgaagttcgaaataattgggcgtatatgtctatcttgac       600
RCA_P40-CabLCV-Clone2      ctcggtgttaaacctttaatttgaagttcgaaataattgggcgtatatgtctatcttgac       600
RCA_P40-CabLCV-Clone3      ctcggtgttaaacctttaatttgaagttcgaaataattgggcgtatatgtctatcttgac       600
RCA_P40-CabLCV-Clone4      ctcggtgttaaacctttaatttgaagttcgaaataattgggcgtatatgtctatcttgac       600
RCA_P40-CabLCV-Clone5      ctcggtgttaaacctttaatttgaagttcgaaataattgggcgtatatgtctatcttgac       600
RCA_P40-CabLCV-Clone6      ctcggtgttaaacctttaatttgaagttcgaaataattgggcgtatatgtctatcttgac       600
                           ************************************************************

DNA_B-CabLCV[U65530]       cggtctttgtaacgacaaagcttctgacacattgtacaatatattgaacgtggcccaatt       660
RCA_P40-CabLCV-Clone2      cggtctttgtaacgacaaagcttctgacacattgtacaatatattgaacgtggcccaatt       660
RCA_P40-CabLCV-Clone3      cggtctttgtaacgacaaagcttctgacacattgtacaatatattgaacgtggcccaatt       660
RCA_P40-CabLCV-Clone4      cggtctttgtaacgacaaagcttctgacacattgtacaatatattgaacgtggcccaatt       660
RCA_P40-CabLCV-Clone5      cggtctttgtaacgacaaagcttctgtcacattgtacaatatattgaacgtggcccaatt       660
RCA_P40-CabLCV-Clone6      cggtctttgtaacgacaaagcttctgacacattgtacaatatattgaacgtggcccaatt       660
                           ************************** *********************************

DNA_B-CabLCV[U65530]       atatttcctctacggagttaggctatcgcttatttgtatttaccttgtctatataatgga       720
RCA_P40-CabLCV-Clone2      atatttcctctacggagttaggctatcgcttatttgtatttaccttgtctatataatgga       720
RCA_P40-CabLCV-Clone3      atatttcctctacggagttaggctatcgcttatttgtatttaccttgtctatataatgga       720
RCA_P40-CabLCV-Clone4      atatttcctctacggagttaggctatcgcttatttgtatttaccttgtctatataatgga       720
RCA_P40-CabLCV-Clone5      atatttcctctacggagttaggctatcgcttatttgtatttaccttgtctatataatgga       720
RCA_P40-CabLCV-Clone6      atatttcctctacggagttaggctatcgcttatttgtatttaccttgtctatataatgga       720
                           ************************************************************

DNA_B-CabLCV[U65530]       cgatagtttaatattttaaatcgtctttgacaataacattaaaataacaaatcattttgt       780
RCA_P40-CabLCV-Clone2      cgatagtttaatattttaaatcgtctttgacaataacattaaaataacaaatcattttgt       780
RCA_P40-CabLCV-Clone3      cgatagtttaatattttaaatcgtctttgacaataacattaaaataacaaatcattttgt       780
RCA_P40-CabLCV-Clone4      cgatagtttaatattttaaatcgtctttgacaataacattaaaataacaaatcattttgt       780
RCA_P40-CabLCV-Clone5      cgatagtttaatattttaaatcgtctttgacaataacattaaaataacaaatcattttgt       780
RCA_P40-CabLCV-Clone6      cgatagtttaatattttaaatcgtctttgacaataacattaaaataacaaatcattttgt       780
                           ************************************************************

DNA_B-CabLCV[U65530]       agaggtgctgagaataatgtatcctacaaagtttaggcgtggggtatcttactctcaaag       840
RCA_P40-CabLCV-Clone2      agaggtgctgagaataatgtatcctacaaagtttaggcgtggggtatcttactctcaaag       840
RCA_P40-CabLCV-Clone3      agaggtgctgagaataatgtatcctacaaagtttaggcgtggggtatcttactctcaaag       840
RCA_P40-CabLCV-Clone4      agaggtgctgagaataatgtatcctacaaagtttaggcgtggggtatcttactctcaaag       840
RCA_P40-CabLCV-Clone5      agaggtgctgagaataatgtatcctacaaagtttaggcgtggggtatcttactctcaaag       840
RCA_P40-CabLCV-Clone6      agaggtgctgagaataatgtatcctacaaagtttaggcgtggggtatcttactctcaaag       840
                           ************************************************************

DNA_B-CabLCV[U65530]       acgatttgtttcacgtaatcaatcgtctaagcgtggaacttttgttagacgcactgatgg       900
RCA_P40-CabLCV-Clone2      acgatttgtttcacgtaatcaatcgtctaagcgtggaacttttgttagacgcactgatgg       900
RCA_P40-CabLCV-Clone3      acgatttgtttcacgtaatcaatcgtctaagcgtggaacttttgttagacgcactgatgg       900
RCA_P40-CabLCV-Clone4      acgatttgtttcacgtaatcaatcgtctaagcgtggaacttttgttagacgcactgatgg       900
RCA_P40-CabLCV-Clone5      acgatttgtttcacgtaatcaatcgtctaagcgtggaacttttgttagacgcactgatgg       900
RCA_P40-CabLCV-Clone6      acgatttgtttcacgtaatcaatcgtctaagcgtggaacttttgttagacgcactgatgg       900
                           ************************************************************

DNA_B-CabLCV[U65530]       gaaacgtcgtaaaggcccatcaagtaaagcccatgatgagcctaaaatgaagttgcaacg       960
RCA_P40-CabLCV-Clone2      gaaacgtcgtaaaggcccatcaagtaaagcccatgatgagcctaaaatgaagttgcaacg       960
RCA_P40-CabLCV-Clone3      gaaacgtcgtaaaggcccatcaagtaaagcccatgatgagcctaaaatgaagttgcaacg       960
RCA_P40-CabLCV-Clone4      gaaacgtcgtaaaggcccatcaagtaaagcccatgatgagcctaaaatgaagttgcaacg       960
RCA_P40-CabLCV-Clone5      gaaacgtcgtaaaggcccatcaagtaaagcccatgatgagcctaaaatgaagttgcaacg       960
RCA_P40-CabLCV-Clone6      gaaacgtcgtaaaggcccatcaagtaaagcccatgatgagcctaaaatgaagttgcaacg       960
                           ************************************************************

DNA_B-CabLCV[U65530]       catacatgaaaatcaatatgggcctgaatttgtcatgacccataactcagccctttcaac       1020
RCA_P40-CabLCV-Clone2      catacatgaaaatcaatatgggcctgaatttgtcatgacccataactcagccctttcaac       1020
RCA_P40-CabLCV-Clone3      catacatgaaaatcaatatgggcctgaatttgtcatgacccataactcagccctttcaac       1020
RCA_P40-CabLCV-Clone4      catacatgaaaatcaatatgggcctgaatttgtcatgacccataactcagccctttcaac       1020
RCA_P40-CabLCV-Clone5      catacatgaaaatcaatatgggcctgaatttgtcatgacccataactcagccctttcaac       1020
RCA_P40-CabLCV-Clone6      catacatgaaaatcaatatgggcctgaatttgtcatgacccataactcagccctttcaac       1020
                           ************************************************************
```

```
DNA_B-CabLCV[U65530]      gtttattaatttccctgtacttggtaagattgaacctaaccgaagcaggtcgtatattaa      1080
RCA_P40-CabLCV-Clone2     gtttattaatttccctgtacttggtaagattgaacctaaccgaagcaggtcgtatattaa      1080
RCA_P40-CabLCV-Clone3     gtttattaatttccctgtacttggtaagattgaacctaaccgaagcaggtcgtatattaa      1080
RCA_P40-CabLCV-Clone4     gtttattaatttccctgtacttggtaagattgaacctaaccgaagcaggtcgtatattaa      1080
RCA_P40-CabLCV-Clone5     gtttattaatttccctgtacttggtaagattgaacctaaccgaagcaggtcgtatattaa      1080
RCA_P40-CabLCV-Clone6     gtttattaatttccctgtacttggtaagattgaacctaaccgaagcaggtcgtatattaa      1080
                          ************************************************************

DNA_B-CabLCV[U65530]      gttgaaccggttatcatttaagggaaccgttaagattgagcgtgtacatgctgatgtgaa      1140
RCA_P40-CabLCV-Clone2     gttgaaccggttatcatttaagggaaccgttaagattgagcgtgtacatgctgatgtgaa      1140
RCA_P40-CabLCV-Clone3     gttgaaccggttatcatttaagggaaccgttaagattgagcgtgtacatgctgatgtgaa      1140
RCA_P40-CabLCV-Clone4     gttgaaccggttatcatttaagggaaccgttaagattgagcgtgtacatgctgatgtgaa      1140
RCA_P40-CabLCV-Clone5     gttgaaccggttatcatttaagggaaccgttaagattgagcgtgtacatgctgatgtgaa      1140
RCA_P40-CabLCV-Clone6     gttgaaccggttatcatttaagggaaccgttaagattgagcgtgtacatgctgatgtgaa      1140
                          ************************************************************

DNA_B-CabLCV[U65530]      catggacggagtaatttcgaagatagagggtgtgttctctcttgttattgttgttgatcg      1200
RCA_P40-CabLCV-Clone2     catggacggagtaatttcgaagatagagggtgtgttctctcttgttattgttgttgatcg      1200
RCA_P40-CabLCV-Clone3     catggacggagtaatttcgaagatagagggtgtgttctctcttgttattgttgttgatcg      1200
RCA_P40-CabLCV-Clone4     catggacggagtaatttcgaagatagagggtgtgttctctcttgttattgttgttgatcg      1200
RCA_P40-CabLCV-Clone5     catggacggagtaatttcgaagatagagggtgtgttctctcttgttattgttgttgatcg      1200
RCA_P40-CabLCV-Clone6     catggacggagtaatttcgaagatagagggtgtgttctctcttgttattgttgttgatcg      1200
                          ************************************************************

DNA_B-CabLCV[U65530]      caaaccacatttaagctccactggaggtttgcatacatttgatgaaatatttggtgcaag      1260
RCA_P40-CabLCV-Clone2     caaaccacatttaagctccactggaggtttgcatacatttgatgaaatatttggtgcaag      1260
RCA_P40-CabLCV-Clone3     caaaccacatttaagctccactggaggtttgcatacatttgatgaaatatttggtgcaag      1260
RCA_P40-CabLCV-Clone4     caaaccacatttaagctccactggaggtttgcatacatttgatgaaatatttggtgcaag      1260
RCA_P40-CabLCV-Clone5     caaaccacatttaagctccactggaggtttgcatacatttgatgaaatatttggtgcaag      1260
RCA_P40-CabLCV-Clone6     caaaccacatttaagctccactggaggtttgcatacatttgatgaaatatttggtgcaag      1260
                          ************************************************************

DNA_B-CabLCV[U65530]      aatccatagccatgggaacttagctattacacccggtttgaaagatcgttattacgtcct      1320
RCA_P40-CabLCV-Clone2     aatccatagccatgggaacttagctattacacccggtttgaaagatcgttattacgtcct      1320
RCA_P40-CabLCV-Clone3     aatccatagccatgggaacttagctattacacccggtttgaaagatcgttattacgtcct      1320
RCA_P40-CabLCV-Clone4     aatccatagccatgggaacttagctattacacccggtttgaaagatcgttattacgtcct      1320
RCA_P40-CabLCV-Clone5     aatccatagccatgggaacttagctattacacccggtttgaaagatcgttattacgtcct      1320
RCA_P40-CabLCV-Clone6     aatccatagccatgggaacttagctattacacccggtttgaaagatcgttattacgtcct      1320
                          ************************************************************

DNA_B-CabLCV[U65530]      ccatgttttgaaacgcgtattgtctgtggagaaagacactttgatggtggatcttgaagg      1380
RCA_P40-CabLCV-Clone2     ccatgttttgaaacgcgtattgtctgtggagaaagacactttgatggtggatcttgaagg      1380
RCA_P40-CabLCV-Clone3     ccatgttttgaaacgcgtattgtctgtggagaaagacactttgatggtggatcttgaagg      1380
RCA_P40-CabLCV-Clone4     ccatgttttgaaacgcgtattgtctgtggagaaagacactttgatggtggatcttgaagg      1380
RCA_P40-CabLCV-Clone5     ccatgttttgaaacgcgtattgtctgtggagaaagacactttgatggtggatcttgaagg      1380
RCA_P40-CabLCV-Clone6     ccatgttttgaaacgcgtattgtctgtggagaaagacact-tgatggtggatcttgaagg      1379
                          **************************************** *******************

DNA_B-CabLCV[U65530]      atctaccacgatatctaataggcgttataactgttgggcctcatttaacgatcttgaaca      1440
RCA_P40-CabLCV-Clone2     atctaccacgatatctaataggcgttataactgttgggcctcatttaacgatcttgaaca      1440
RCA_P40-CabLCV-Clone3     atctaccacgatatctaataggcgttataactgttgggcctcatttaacgatcttgaaca      1440
RCA_P40-CabLCV-Clone4     atctaccacgatatctaataggcgttataactgttgggcctcatttaacgatcttgaaca      1440
RCA_P40-CabLCV-Clone5     atctaccacgatatctaataggcgttataactgttgggcctcatttaacgatcttgaaca      1440
RCA_P40-CabLCV-Clone6     atctaccacgatatctaataggcgttataactgttgggcctcatttaacgatcttgaaca      1439
                          ************************************************************

DNA_B-CabLCV[U65530]      tgacttatgtaacggtgtttatgcgaatataagcaaaaacgccattttggtatattattg      1500
RCA_P40-CabLCV-Clone2     tgacttatgtaacggtgtttatgcgaatataagcaaaaacgccattttggtatattattg      1500
RCA_P40-CabLCV-Clone3     tgacttatgtaacggtgtttatgcgaatataagcaaaaacgccattttggtatattattg      1500
RCA_P40-CabLCV-Clone4     tgacttatgtaacggtgtttatgcgaatataagcaaaaacgccattttggtatattattg      1500
RCA_P40-CabLCV-Clone5     tgacttatgtaacggtgtttatgcgaatataagcaaaaacgccattttggtatattattg      1500
RCA_P40-CabLCV-Clone6     tgacttatgtaacggtgtttatgcgaatataagcaaaaacgccattttggtatattattg      1499
                          ************************************************************

DNA_B-CabLCV[U65530]      ttggatgtccgatgctatgtctaaggcatcgacctttgtatcttacgatcttgattattt      1560
RCA_P40-CabLCV-Clone2     ttggatgtccgatgctatgtctaaggcatcgacctttgtatcttacgatcttgattattt      1560
RCA_P40-CabLCV-Clone3     ttggatgtccgatgctatgtctaaggcatcgacctttgtatcttacgatcttgattattt      1560
RCA_P40-CabLCV-Clone4     ttggatgtccgatgctatgtctaaggcatcgacctttgtatcttacgatcttgattattt      1560
RCA_P40-CabLCV-Clone5     ttggatgtccgatgctatgtctaaggcatcgacctttgtatcttacgatcttgattattt      1560
RCA_P40-CabLCV-Clone6     ttggatgtccgatgctatgtctaaggcatcgacctttgtatcttacgatcttgattattt      1559
                          ************************************************************

DNA_B-CabLCV[U65530]      aggttaaccatgaataaaatggcgttaagattgaatatatgcattaataataaaacacgc      1620
RCA_P40-CabLCV-Clone2     aggttaaccatgaataaaatggcgttaagattgaatatatgcattaataataaaacacgc      1620
RCA_P40-CabLCV-Clone3     aggttaaccatgaataaaatggcgttaagattgaatatatgcattaataataaaacacgc      1620
RCA_P40-CabLCV-Clone4     aggttaaccatgaataaaatggcgttaagattgaatatatgcattaataataaaacacgc      1620
RCA_P40-CabLCV-Clone5     aggttaaccatgaataaaatggcgttaagattgaatatatgcattaataataaaacacgc      1620
RCA_P40-CabLCV-Clone6     aggttaaccatgaataaaatggcgttaagattgaatatatgcattaataataaaacacgc      1619
                          ************************************************************
```

```
DNA_B-CabLCV[U65530]      aattttattgcaatgactttggttgtggaggattacaattattgttaatacattcctgaa      1680
RCA_P40-CabLCV-Clone2     aattttattgcaatgactttggttgtggaggattacaattattgttaatacattcctgaa      1680
RCA_P40-CabLCV-Clone3     aattttattgcaatgactttggttgtggaggattacaattattgttaatacattcctgaa      1680
RCA_P40-CabLCV-Clone4     aattttattgcaatgactttggttgtggaggattacaattattgttaatacattcctgaa      1680
RCA_P40-CabLCV-Clone5     aattttattgcaatgactttggttgtggaggattacaattattgttaatacattcctgaa      1680
RCA_P40-CabLCV-Clone6     aattttattgcaatgactttggttgtggaggattacaattattgttaatacattcctgaa      1679
                          ************************************************************

DNA_B-CabLCV[U65530]      ccgtcgtcctaactagctcgcttaattgggccactgacatcgttatggttgattgggccc      1740
RCA_P40-CabLCV-Clone2     ccgtcgtcctaactagctcgcttaattgggccactgacatcgttatggttgattgggccc      1740
RCA_P40-CabLCV-Clone3     ccgtcgtcctaactagctcgcttaattgggccactgacatcgttatggttgattgggccc      1740
RCA_P40-CabLCV-Clone4     ccgtcgtcctaactagctcgcttaattgggccactgacatcgttatggttgattgggccc      1740
RCA_P40-CabLCV-Clone5     ccgtcgtcctaactagctcgcttaattgggccactgacatcgttatggttgattgggccc      1740
RCA_P40-CabLCV-Clone6     ccgtcgtcctaactagctcgcttaattgggccactgacatcgttatggttgattgggccc      1739
                          ************************************************************

DNA_B-CabLCV[U65530]      tttgtaacccagcttgtgatgctgaatccccggggtctaatgcgctagttccta-gcagg      1799
RCA_P40-CabLCV-Clone2     tttgtaacccagcttgtgatgctgaatccccggggtctaatgcgctagttccta-gcagg      1799
RCA_P40-CabLCV-Clone3     tttgtaacccagcttgtgatgctgaatccccggggtctaatgcgctagttcctaagcagg      1800
RCA_P40-CabLCV-Clone4     tttgtaacccagcttgtgatgctgaatccccggggtctaatgcgctagttccta-gcagg      1799
RCA_P40-CabLCV-Clone5     tttgtaacccagcttgtgatgctgaatccccggggtctaatgcgctagttccta-gcagg      1799
RCA_P40-CabLCV-Clone6     tttgtaacccagcttgtgatgctgaatccccggggtctaatgcgctagttccta-gcagg      1798
                          ****************************************************** *****

DNA_B-CabLCV[U65530]      ttgagttctctatatggatgtagcgcgttttccacttctgattctgtgtgtgggttggga      1859
RCA_P40-CabLCV-Clone2     ttgagttctctatatggatgtagcgcgttttccacttctgattctgtgtgtgggttggga      1859
RCA_P40-CabLCV-Clone3     ttgagttctctatatggatgtagcgcgttttccacttctgattctgtgtgtgggttggga      1860
RCA_P40-CabLCV-Clone4     ttgagttctctatatggatgtagcgcgttttccacttctgattctgtgtgtgggttggga      1859
RCA_P40-CabLCV-Clone5     ttgagttctctatatggatgtagcgcgttttccacttctgattctgtgtgtgggttggga      1859
RCA_P40-CabLCV-Clone6     ttgagttctctatatggatgtagcgcgttttccacttctgattcg-ggtgtgggttggga      1857
                          ********************************************   *************

DNA_B-CabLCV[U65530]      aacccaattgtgctccttgaagcccatgaatcacctggttgtaattcaattgggcctggg      1919
RCA_P40-CabLCV-Clone2     aacccaattgtgctccttgaagcccatgaatcacctggttgtaattcaattgggcctggg      1919
RCA_P40-CabLCV-Clone3     aacccaattgtgctccttgaagcccatgaatcacctggttgtaattcaattgggcctggg      1920
RCA_P40-CabLCV-Clone4     aacccaattgtgctccttgaagcccatgaatcacctggttgtaattcaattgggcctggg      1919
RCA_P40-CabLCV-Clone5     aacccaattgtgctccttgaagcccatgaatcacctggttgtaattcaattgggcctggg      1919
RCA_P40-CabLCV-Clone6     aacccaattgtgctccttgaagcccctgaatcacctggttgtaattcaattgggcctggg      1917
                          ************************* **********************************

DNA_B-CabLCV[U65530]      cctgttagtccaattcttgacaatgatttggacctcaaggtcttcctctcccatcttccg      1979
RCA_P40-CabLCV-Clone2     cctgttagtccaattcttgacaatgatttggacctcaaggtcttcctctcccatcttccg      1979
RCA_P40-CabLCV-Clone3     cctgttagtccaattcttgacaatgatttggacctcaaggtcttcctctcccatcttccg      1980
RCA_P40-CabLCV-Clone4     cctgttagtccaattcttgacaatgatttggacctcaaggtcttcctctcccatcttccg      1979
RCA_P40-CabLCV-Clone5     cctgttagtccaattcttgacaatgatttggacctcaaggtcttcctctcccatcttccg      1979
RCA_P40-CabLCV-Clone6     cctgttagtccaattcttgacaatgatttggacctcaaggtcttcctctcccatcttccg      1977
                          ************************************************************

DNA_B-CabLCV[U65530]      tagtccacatgtgagaaatcaacatctctatgtgaaaattgttttgaatgaattttcact      2039
RCA_P40-CabLCV-Clone2     tagtccacatgtgagaaatcaacatctctatgtgaaaattgttttgaatgaattttcact      2039
RCA_P40-CabLCV-Clone3     tagtccacatgtgagaaatcaacatctctatgtgaaaattgttttgaatgaattttcact      2040
RCA_P40-CabLCV-Clone4     tagtccacatgtgagaaatcaacatctctatgtgaaaattgttttgaatgaattttcact      2039
RCA_P40-CabLCV-Clone5     tagtccacatgtgagaaatcaacatctctatgtgaaaattgttttgaatgaattttcact      2039
RCA_P40-CabLCV-Clone6     tagtccacatgtgagaaatcaacatctctatgtgaaaattgttttgaatgaattttcact      2037
                          ************************************************************

DNA_B-CabLCV[U65530]      gttggtgcccggaaggggatatccactgaatgtttagctgtggacaatttcaatttcccc      2099
RCA_P40-CabLCV-Clone2     gttggtgcccggaaggggatatccactgaatgtttagctgtggacaatttcaatttcccc      2099
RCA_P40-CabLCV-Clone3     gttggtgcccggaaggggatatccactgaatgtttagctgtggacaatttcaatttcccc      2100
RCA_P40-CabLCV-Clone4     gttggtgcccggaaggggatatccactgaatgtttagctgtggacaatttcaatttcccc      2099
RCA_P40-CabLCV-Clone5     gttggtgcccggaaggggatatccactgaatgtttagctgtggacaatttcaatttcccc      2099
RCA_P40-CabLCV-Clone6     gttggtgcccggaaggggatatccactgaatgtttagctgtggacaatttcaatttcccc      2097
                          ************************************************************

DNA_B-CabLCV[U65530]      ttaaacttggcaaaatgtgttcgttgatgcacatttgtgtcgctaactctgtaatagagt      2159
RCA_P40-CabLCV-Clone2     ttaaacttggcaaaatgtgttcgttgatgcacatttgtgtcgctaactctgtaatagagt      2159
RCA_P40-CabLCV-Clone3     ttaaacttggcaaaatgtgttcgttgatgcacatttgtgtcgctaactctgtaatagagt      2160
RCA_P40-CabLCV-Clone4     ttaaacttggcaaaatgtgttcgttgatgcacatttgtgtcgctaactctgtaatagagt      2159
RCA_P40-CabLCV-Clone5     ttaaacttggcaaaatgtgttcgttgatgcacatttgtgtcgctaactctgtaatagagt      2159
RCA_P40-CabLCV-Clone6     ttaaacttggcaaaatgtgttcgttgatgcacatttgtgtcgctaactctgtaatagagt      2157
                          ************************************************************

DNA_B-CabLCV[U65530]      ttccacggaatggggtcttttagagagaagaatgaagatgaaaaatagtggagatctatg      2219
RCA_P40-CabLCV-Clone2     ttccacggaatggggtcttttagagagaagaatgaagatgaaaaatagtggagatctatg      2219
RCA_P40-CabLCV-Clone3     ttccacggaatggggtcttttagagagaagaatgaagatgaaaaatagtggagatctatg      2220
RCA_P40-CabLCV-Clone4     ttccacggaatggggtcttttagagagaagaatgaagatgaaaaatagtggagatctatg      2219
RCA_P40-CabLCV-Clone5     ttccacggaatggggtcttttagagagaagaatgaagatgaaaaatagtggagatctatg      2219
RCA_P40-CabLCV-Clone6     ttccacggaatggggtcttttagagagaagaatgaagatgaaaaatagtggagatctatg      2217
                          ************************************************************
```

```
DNA_B-CabLCV[U65530]        ttgcatcttaagggaaatgtccaagacgcttgtaaggattcgtcgtcagtcatcctcttg        2279
RCA_P40-CabLCV-Clone2       ttgcatcttaagggaaatgtccaagacgcttgtaaggattcgtcgtcagtcatcctcttg        2279
RCA_P40-CabLCV-Clone3       ttgcatcttaagggaaatgtccaagacgcttgtaaggattcgtcgtcagtcatcctcttg        2280
RCA_P40-CabLCV-Clone4       ttgcatcttaagggaaatgtccaagacgcttgtaaggattcgtcgtcagtcatcctcttg        2279
RCA_P40-CabLCV-Clone5       ttgcatcttaagggaaatgtccaagacgcttgtaaggattcgtcgtcagtcatcctcttg        2279
RCA_P40-CabLCV-Clone6       ttgcatcttaagggaaatgtccaagacgcttgtaaggattcgtcgtcagtcatcctcttg        2277
                            ************************************************************

DNA_B-CabLCV[U65530]        tcatggatctccactatcacagatccagttgcgttaattggtacttgctgtctatattcg        2339
RCA_P40-CabLCV-Clone2       tcatggatctccactatcacagatccagttgcgttaattggtacttgctgtctatattcg        2339
RCA_P40-CabLCV-Clone3       tcatggatctccactatcacagatccagttgcgttaattggtacttgctgtctatattcg        2340
RCA_P40-CabLCV-Clone4       tcatggatctccactatcacagatccagttgcgttaattggtacttgctgtctatattcg        2339
RCA_P40-CabLCV-Clone5       tcatggatctccactatcacagatccagttgcgttaattggtacttgctgtctatattcg        2339
RCA_P40-CabLCV-Clone6       tcatggatctccactatcacagatccagttgcgttaattggtacttgctgtctatattcg        2337
                            ************************************************************

DNA_B-CabLCV[U65530]        atgacgcaatggtcgattttcatacagctacgattaagccttgctgtaaattgcgctgca        2399
RCA_P40-CabLCV-Clone2       atgacgcaatggtcgattttcatacagctacgattaagccttgctgtaaattgcgctgca        2399
RCA_P40-CabLCV-Clone3       atgacgcaatggtcgattttcatacagctacgattaagccttgctgtaaattgcgctgca        2400
RCA_P40-CabLCV-Clone4       atgacgcaatggtcgattttcatacagctacgattaagccttgctgtaaattgcgctgca        2399
RCA_P40-CabLCV-Clone5       atgacgcaatggtcgattttcatacagctacgattaagccttgctgtaaattgcgctgca        2399
RCA_P40-CabLCV-Clone6       atgacgcaatggtcgattttcatacagctacgattaagccttgctgtaaattgcgctgca        2397
                            ************************************************************

DNA_B-CabLCV[U65530]        gttgaagggaattgaagtagtatctcagtaagatcatgcgacagctgatattcgtctcta        2459
RCA_P40-CabLCV-Clone2       gttgaagggaattgaagtagtatctcagtaagatcatgcgacagctgatattcgtctcta        2459
RCA_P40-CabLCV-Clone3       gttgaagggaattgaagtagtatctcagtaagatcatgcgacagctgatattcgtctcta        2460
RCA_P40-CabLCV-Clone4       gttgaagggaattgaagtagtatctcagtaagatcatgcgacagctgatattcgtctcta        2459
RCA_P40-CabLCV-Clone5       gttgaagggaattgaagtagtatctcagtaagatcatgcgacagctgatattcgtctcta        2459
RCA_P40-CabLCV-Clone6       gttgaagggaattgaagtagtatctcagtaagatcatgcgacagctgatattcgtctcta        2457
                            ************************************************************

DNA_B-CabLCV[U65530]        tgagactctatgtaattaaacgcatttggagcattcgctaactgagaattcat       2512
RCA_P40-CabLCV-Clone2       tgagactctatgtaattaaacgcatttggagcattcgctaactgagaattcat       2512
RCA_P40-CabLCV-Clone3       tgagactctatgtaattaaacgcatttggagcattcgctaactgagaattcat       2513
RCA_P40-CabLCV-Clone4       tgagactctatgtaattaaacgcatttggagcattcgctaactgagaattcat       2512
RCA_P40-CabLCV-Clone5       tgagactctatgtaattaaacgcatttggagcattcgctaactgagaattcat       2512
RCA_P40-CabLCV-Clone6       tgagactctatgtaattaaacgcatttggagcattcgctaactgagaattcat       2510
                            *****************************************************
```

**Fig S3.** Multiple Sequence Alignment of consensus CabLCV sequences generated from Sanger sequence of 6 independent RCA-generated clones. Both CabLCV DNA A (1/6) and DNA B (5/6) genomic components were detected in P40 fractions. Alignments were performed using the ClustalW alignment tool from the European Bioinformatics Institute (EBI, ebi.ac.uk).

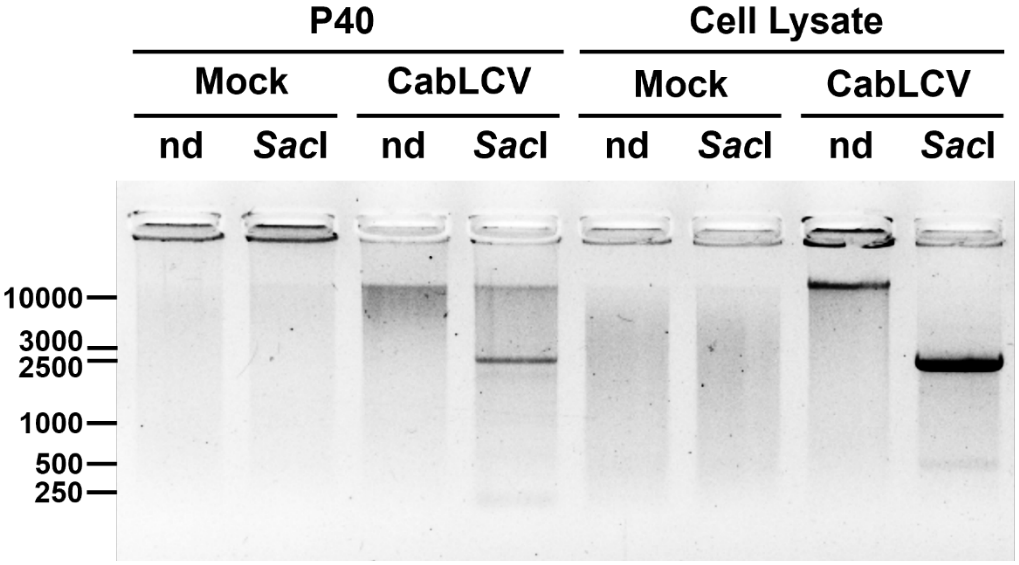


**Fig. S4.** P40 fractions from from CabLCV-infected *Nicotiana benthamiana* plants contain complete geminiviral genomes. Rolling Circle Amplification (RCA) allowed detection of complete circular CabLCV genomes in P40 fractions from infected *N. benthamiana* plants. RCA products were linearized using a single-cutter endonuclease (*Sac*I). RCA using cell lysate DNA extractions as templates were included as positive controls. **nd**: non-digested. ***Sac*I**: *Sac*I-digested.

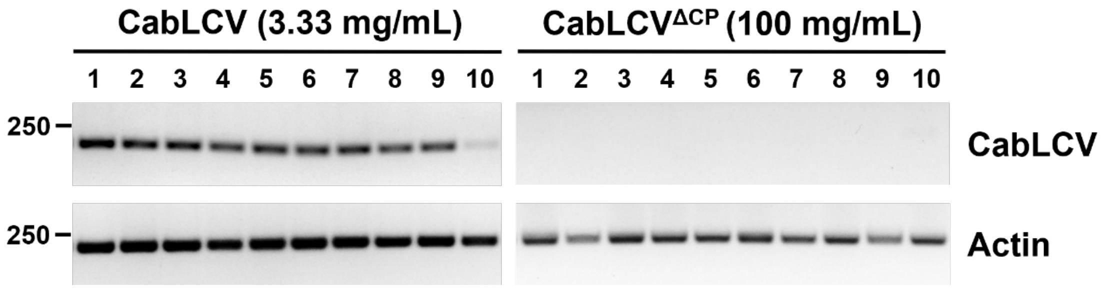


**Fig. S5.** No viral DNA amplification was achieved when Arabidopsis thaliana plants were mechanically inoculated with extracts from CabLCV$^{\Delta CP}$-infected plants (right). Symptomatic *A. thaliana* plants inoculated with equivalent doses of DNA A from wild-type CabLCV were included as controls (left). Experiment was repeated twice with similar results.

### Tables

**Table S1. Relative densitometry analysis of EV-marker protein increase after immunodetection.** Similar results were obtained in two independent biological replicates.

| | R1 | | R2 | | Mean Ratio |
|---|---|---|---|---|---|
| | **Mock** | **CabLCV** | **Mock** | **CabLCV** | **CabLCV/Mock** |
| **RFP-PEN1** | 1 | 1.92 | 1 | 3.66 | 2.79 |
| **TET8-GFP** | 1 | 1.47 | 1 | 2.02 | 1.74 |
| **PATL1** | 1 | 3.90 | 1 | 1.46 | 2.68 |

**Table S2. CabLCV P40 proteome is enriched in apoplast-related GO categories.** Main apoplast-related categories detected in the CabLCV-infected P40 proteome in a Gene Ontology Cellular Component (GO-CC) functional enrichment (STRING; string-db.org).

| #term ID | Term description | Observed gene count | Background gene count | Strength | Signal | False discovery rate (FDR) |
|---|---|---|---|---|---|---|
| GO:0099503 | Secretory vesicle | 124 | 165 | 1.02 | 5.62 | 2.04E-64 |
| GO:0048046 | Apoplast | 208 | 450 | 0.81 | 4.14 | 5.35E-81 |
| GO:0009505 | Plant-type cell wall | 169 | 368 | 0.08 | 3.94 | 1.06E-64 |
| GO:0005618 | Cell wall | 175 | 402 | 0.78 | 3.77 | 2.12E-64 |
| GO:0030312 | External encapsulating structure | 180 | 429 | 0.77 | 3.67 | 2.12E-64 |
| GO:0031982 | Vesicle | 205 | 885 | 0.51 | 2.07 | 3.82E-39 |

**Table S3. CabLCV$^{\Delta CP}$ DNA accumulation in Arabidopsis is significantly lower compared to wild-type CabLCV.** CabLCV DNA viral accumulation was measured by qPCR from three pools (20 plants each) of CabLCV WT and mutant CabLCV$^{\Delta CP}$ infected plants at 21 dpi. Values were normalized and calibrated to 1 for the CabLCV WT viral accumulation and represent mean and standard error.

| Relative viral DNA accumulation | DNA A | DNA B |
|---|---|---|
| **CabLCV** | 1 ± 0.06 | 1 ± 0.00 |
| **CabLCV$^{\Delta CP}$** | 0.04 ± 0.02 | 0.11 ± 0.01 |

**Table S4.** List of antibodies used in this work.

| Antibody | Species | Dilution | Source | Catalog number |
|---|---|---|---|---|
| RED FLUORESCENT PROTEIN (RFP) | Mouse | 1:1000 | Chromotek | 6G6 |
| GREEN FLUORESCENT PROTEIN (GFP) | Mouse | 1:2000 | Sigma-Aldrich | 66002-1-Ig |
| PATELLIN 1 (PATL1) | Rabbit | 1:5000 | Peterman et al., 2004 | N/A |
| SYNTAXIN 61 (SYP61) | Rabbit | 1:1000 | Sanderfoot et al., 2001, ABRC | AB00125 |
| Rabbit-HRP | Goat | 1:10000 | Abcam | AB97051 |
| Mouse-HRP | Goat | 1:10000 | Abcam | AB6789 |

**Table S5.** List of oligonucleotides used in this work.

| Primer Name | Sequence | Target gene | Use | Reference |
|---|---|---|---|---|
| CabLCV AL3 Fw<br>CabLCV AL3 Rv | AATATGTCGGCCCAGAAGC<br>CAGGATACAACGCGTAGAGGA | *AL3* from DNA-A of Cabbage leaf curl virus (U65529) | PCR detection of CabLCV DNA-A and sequencing | Mauricio-Zúñiga, unpublished |
| CabLCV BC1 Fw<br>CabLCV BC1 Rv | GTCCTAACTAGCTCGCTTAATTGGG<br>CGCGCTACATCCATATAGAGAACTC | *BC1* from DNA-B of Cabbage leaf curl virus (U65530) | PCR detection of CabLCV DNA-B | This work |
| upV2BC-qRT<br>lowV2BC-qRT | ATGGGACCTTTCAGAGTGGA<br>GACGAAAGACCTCGCCTTCT | *V2* from Beet curly top virus – California [Logan] (NC001412) | PCR detection of BCTV | Luna et al. (2017) |
| AtActin Fw<br>AtActin Rv | CTAAGCTCTCAAGATCAAAGGCTTA<br>ACTAAAACGCAAAACGAAAGCGGTT | *ACT2* from *Arabidopsis thaliana* (*AT3G18780*) | Amplification of actin from *Arabidopsis thaliana* | McKinney & Meagher (1998) |
| Universal M13 Fw<br>Universal M13 Rv | TGTAAAACGGCCAGT<br>CAGGAAACAGCTATGAC | Sequencing from M13 sites | Sanger sequencing of CabLCV RCA clones from P40 | Universal primers |
| CabLCV BC1 Seq Fw<br>CabLCV BC1 Seq Rv | TTGGGCCACTGACATCGTTATG<br>TTGGGTTTCCCAACCCACACAC | *BC1* from DNA-B of Cabbage leaf curl virus (U65530) | Sanger sequencing of CabLCV DNA B | Pedraza-Rubio, unpublished |

**Dataset S1 (separate file).** CabLCV-infected P40 proteome.
